# OmicClaw: executable and reproducible natural-language multi-omics analysis over the unified OmicVerse ecosystem

**DOI:** 10.64898/2026.03.13.711464

**Authors:** Zehua Zeng, Xuehai Wang, Zhi Luo, Yawen Zheng, Lei Hu, Cencan Xing, Hongwu Du

## Abstract

Advances in bulk, single-cell and spatial omics have transformed biological discovery, yet analysis remains fragmented across packages with incompatible interfaces, heterogeneous dependencies and limited workflow reproducibility. Here, we present OmicClaw, an executable natural-language framework for multi-omics analysis built on the unified OmicVerse ecosystem and the J.A.R.V.I.S. runtime. OmicVerse organizes upstream processing, preprocessing, single-cell, spatial, bulk-transcriptomic and foundation-model workflows into a shared AnnData-centered interface spanning more than 100 methods. J.A.R.V.I.S. converts this ecosystem into a bounded analytical action space through a registry-grounded, state-aware and recoverable execution layer that validates prerequisites, preserves provenance and supports iterative repair, while enabling conversational, notebook and visual interfaces to operate over the same live analytical state. Rather than relying on unconstrained code generation, OmicClaw translates user requests into traceable workflows over live omics objects. Across a benchmark of 15 tasks spanning scRNA-seq, spatial transcriptomics, RNA velocity, scATAC-seq, CITE-seq and multiome analysis, ov.Agent improved rubric-based performance over bare one-shot large language model baselines, particularly for long-horizon multi-step workflows. OmicClaw further supports external agent access through an MCP-compatible server and a beginner-friendly web platform for interactive analysis, code execution and million-scale visualization. Together, OmicClaw provides a practical foundation for reproducible human–AI collaboration in modern multi-omics research

---

Recent advances in bulk, single-cell and spatial omics have greatly expanded the scope of biological discovery, but they have also exposed a persistent software bottleneck^12^. Core analytical tasks including preprocessing, clustering, cell-type annotation, trajectory inference, RNA velocity, cell–cell communication, gene regulatory network analysis and spatial deconvolution, are often distributed across independent packages with incompatible interfaces, inconsistent object conventions, and heterogeneous dependencies^3–5^. In practice, users must repeatedly bridge tools that were not designed to interoperate, write wrappers around method-specific APIs and manually reorganize intermediate outputs across workflows. This fragmentation increases the technical burden, weakens reproducibility and makes it difficult to build robust multi-step pipelines that span analytical tasks and omics modalities.

Here, we present **OmicClaw**, an integrated framework for executable natural-language multi-omics analysis built on Omicverse and **J.A.R.V.I.S. (Fig.1a)**. Within this framework, **OmicVerse** provides the analytical substrate: an AnnData-centered ecosystem that organizes diverse omics methods into task-oriented modules and shared interfaces (**Supplementary Table 1**). **J.A.R.V.I.S.** provides the execution layer: a registry-grounded runtime that translates user requests into validated executable workflows over the OmicVerse function space while preserving state, provenance and recovery. Together, these layers extend OmicVerse beyond a collection of individual methods into a coherent analytical environment for bulk, single-cell, and spatial omics. This design differs from ecosystem-level infrastructures such as scverse^4^, domain-specific packages such as Scanpy^6^, and general-purpose bioinformatics toolkits such as Biopython^7^ by emphasizing both interface harmonization across analytical tasks and execution through a bounded, inspectable function space. We therefore do not argue that an LLM alone can perform bioinformatics well; rather, we show that natural-language analysis becomes substantially more executable and reproducible when it is grounded in a unified omics ecosystem and carried out through a bounded, inspectable execution runtime.

**Fig. 1.**
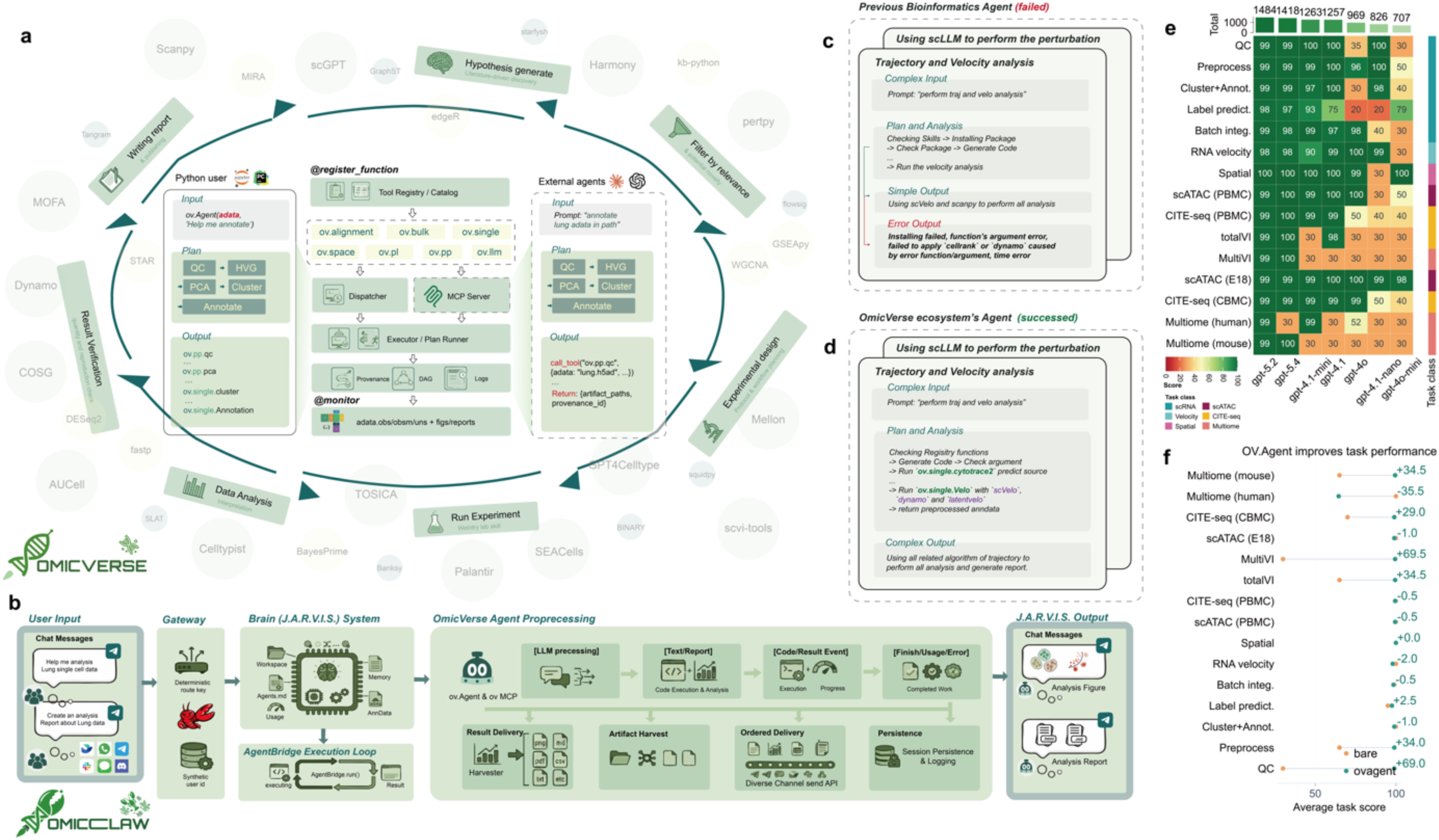
J.A.R.V.I.S. extends OmicVerse into a unified, agentic ecosystem for interoperable multi-omics analysis. **a,** Overview of the OmicClaw ecosystem augmented with **J.A.R.V.I.S.**, an agentic layer that unifies heterogeneous bioinformatics tools through a shared registry and execution framework. Native Python users can directly invoke OmicVerse functions through ov.Agent, whereas third-party or external agent clients can access the same functionality through the MCP server and tool-calling interface. At the core of the system, functions registered by @register_function are organized into a centralized tool registry/catalog spanning multiple OmicVerse modules, including alignment, bulk, single-cell, spatial, visualization, preprocessing and LLM-enabled analysis. Requests are routed through a dispatcher and executor/plan runner, with provenance tracking, directed acyclic graph (DAG) recording and logs enabling transparent monitoring and reproducible execution. The outer loop illustrates the broader closed-loop workflow supported by the ecosystem, including hypothesis generation, relevance filtering, experimental design, experiment execution, data analysis, result verification and report writing. **b,** End-to-end workflow of the J.A.R.V.I.S. system. User requests are first interpreted at the gateway layer, then passed to the J.A.R.V.I.S. “brain”, which maintains workspace context, memory, agent state, usage information and AnnData-aware execution. The OmicVerse agent preprocessing layer coordinates LLM reasoning, code execution, result harvesting, artifact collection, ordered multi-channel delivery and session persistence. Final outputs are returned to the user as chat responses together with derived artifacts, such as analysis figures and reports. **c,** Illustration of a conventional bioinformatics agent that fails to robustly complete a complex single-cell analysis task. Although prompted to perform trajectory and velocity analysis, the agent relies on ad hoc package installation and loosely specified tool usage, resulting in argument errors, failed function calls and incomplete outputs. **d,** In contrast, the OmicVerse ecosystem agent successfully resolves the same task by grounding execution in the registry-defined function space. The agent first checks available registered functions, verifies required arguments and preprocessing steps, and then dispatches standardized OmicVerse interfaces to multiple trajectory and velocity backends (for example, scVelo, dynamo and latentvelo). This registry-guided orchestration enables reliable execution and richer outputs, including preprocessed AnnData objects, analysis results and reports. **e,** Heatmap of benchmark scores for ov.Agent across 15 tasks and seven evaluated foundation models. Each cell shows the rubric-based task score (0–100), integrating execution success, artifact validity, scientific reasonableness, reproducibility, efficiency, and hidden-label accuracy where applicable. Numbers above columns indicate total scores across all tasks; the right annotation marks task class. **f**, Task-wise comparison of average scores between the bare one-shot LLM baseline and ov.Agent. Points show mean task scores, and labels indicate score differences (ov.Agent − bare). ov.Agent consistently outperforms the bare baseline across most tasks, highlighting the value of tool orchestration, iterative repair, and contract-aware execution.

To make natural-language bioinformatics analysis executable and reliable, we designed J.A.R.V.I.S. around a registry-grounded function and <u>Model Context Protocol(MCP)</u> framework that converts the OmicVerse ecosystem into a bounded analytical action space (**Fig. 1a**). Rather than allowing a large language model(LLM) to generate code freely against an unconstrained software environment, OmicVerse functions are exposed through @register_function to a centralized tool registry, in which callable operations, argument schemas, and expected outputs are inspectable before execution (**See Supplementary Methods**). This registry is shared by native Python users, ov.Agent, and external agents through an MCP-compatible interface, ensuring that analysis requests are resolved against the same verified function space rather than against undocumented package behaviors. More than 200 functions or classes are registered in the Omicverse system (**See Supplementary Methods**, **Supplementary Table 2**). In practice, this design addresses two common failure modes in AI-assisted bioinformatics analysis: **code hallucination**^8^, in which agents invent invalid function calls, misuse arguments or assemble non-executable pipelines, and **method selection errors**^8,9^, in which agents recommend methods without grounded access to their actual implementation or input constraints. By routing requests through a registry-aware dispatcher and executor while recording provenance, workflow structure, and execution logs, J.A.R.V.I.S. transforms natural-language intent into traceable function calls over live omics objects, making tool use constrained, inspectable and reproducible rather than speculative.

On top of this registry-grounded function layer, we designed OmicClaw as a state-aware and recoverable runtime for iterative biological data analysis (**Fig.1b**). Rather than treating each request as a one-shot prompt–response event, the system maintains an execution context that couples user intent to the current analytical state, including the active AnnData object, prior session history, recent failures, working context and generated artifacts. This design allows user goals to evolve across multiple turns while preserving analytical continuity. The runtime separates model reasoning from action execution, artifact collection and ordered result delivery, enabling the system to return not only conversational responses but also structured outputs such as figures, tables, reports and session records. It further includes plateau detection, prerequisite-aware control logic, domain-specific code transformation and multi-stage recovery from common execution failures. As a result, natural-language interaction is converted from informal analysis suggestion into bounded, interruptible and recoverable workflow execution (**See Supplementary Methods**).

The practical value of this design is illustrated by multi-step analyses such as trajectory and velocity inference. In generic agent settings, a request such as “perform trajectory and velocity analysis” may fail because tools availability is assumed rather than verified, or argument conventions differ across packages, and successful completion depends heavily on the model planning an entire workflow correctly in a single pass (**Fig.1c**). In J.A.R.V.I.S. for OmicVerse, the same request is first grounded in the registry, enabling the system to inspect available skills, validate required arguments and identify prerequisite preprocessing steps before execution. The runtime then executes the task through a long-horizon, state-aware process in which session context, intermediate results and recent failures are continuously incorporated into the next action, and more complex requests can be decomposed into bounded subproblems within the same recoverable execution framework. It can therefore dispatch analysis through standardized OmicVerse interfaces for quality control, dimensionality reduction, clustering, annotation, trajectory inference and velocity-related back ends while preserving a unified output structure and execution trace (**Fig.1d**). This design reduces brittle behavior, improves recovery from execution errors and lowers dependence on raw frontier-model capability alone. In our benchmark, it also narrows the performance gap between general-purpose models and more advanced reasoning models on long-horizon multi-step omics tasks (**Fig.1e-1f**). In addition, before dispatching any tool, the runtime validates that required data structures exist on the AnnData object—for example, confirming the presence of a scaled layer before principal component analysis or a PCA embedding before neighbor-graph construction. If a prerequisite is unmet, the system returns the specific missing requirement together with the suggested next step, preventing the silent failures and invalid execution orderings that frequently arise in LLM-generated pipelines.

The foundation of OmicClaw is **OmicVerse**, components also include optimized implementations for CUDA-enabled back ends and Apple Metal/MPS environments, as well as Python reimplementations of methods previously available mainly in R-based workflows (**Supplementary Table 3**). J.A.R.V.I.S. operates over the OmicVerse ecosystem, which organizes multi-omics analysis into interoperable analytical families. ov.alignment supports conversion of sequencing reads into count matrices for conventional and single-cell sequencing workflows, including velocity-compatible outputs (**Fig.2a**). ov.preprocess standardizes foundational operations such as quality control, highly variable gene selection, dimensionality reduction and batch correction (**Fig.2b**). ov.single unifies clustering, automated cell-type annotation, trajectory inference, RNA velocity and higher-order cell-structure analyses such as cell–cell interaction, differential expression and gene regulatory network inference (**Fig.2c**). ov.space extends the same task-oriented design to spatial omics, including cell segmentation, cross-section alignment, deconvolution and spatial dynamics (**Fig. 2d**). ov.Agent provides a language-driven entry point into this ecosystem by accepting an AnnData object together with a task description, mapping the request onto registry-defined functions through an LLM-mediated interface, and returning executable function calls rather than unconstrained code generation (**Fig. 2e**). ov.bulk supports bulk transcriptomic workflows (**Fig. 2f**), whereas ov.fm provides unified access to biological foundation models (**Fig. 2g**). The key contribution of OmicVerse is therefore not only method breadth, but ecosystem-level interface harmonization, which is also the precondition that makes reliable agent-mediated execution possible. (**Fig.2h**).

**Fig. 2.**
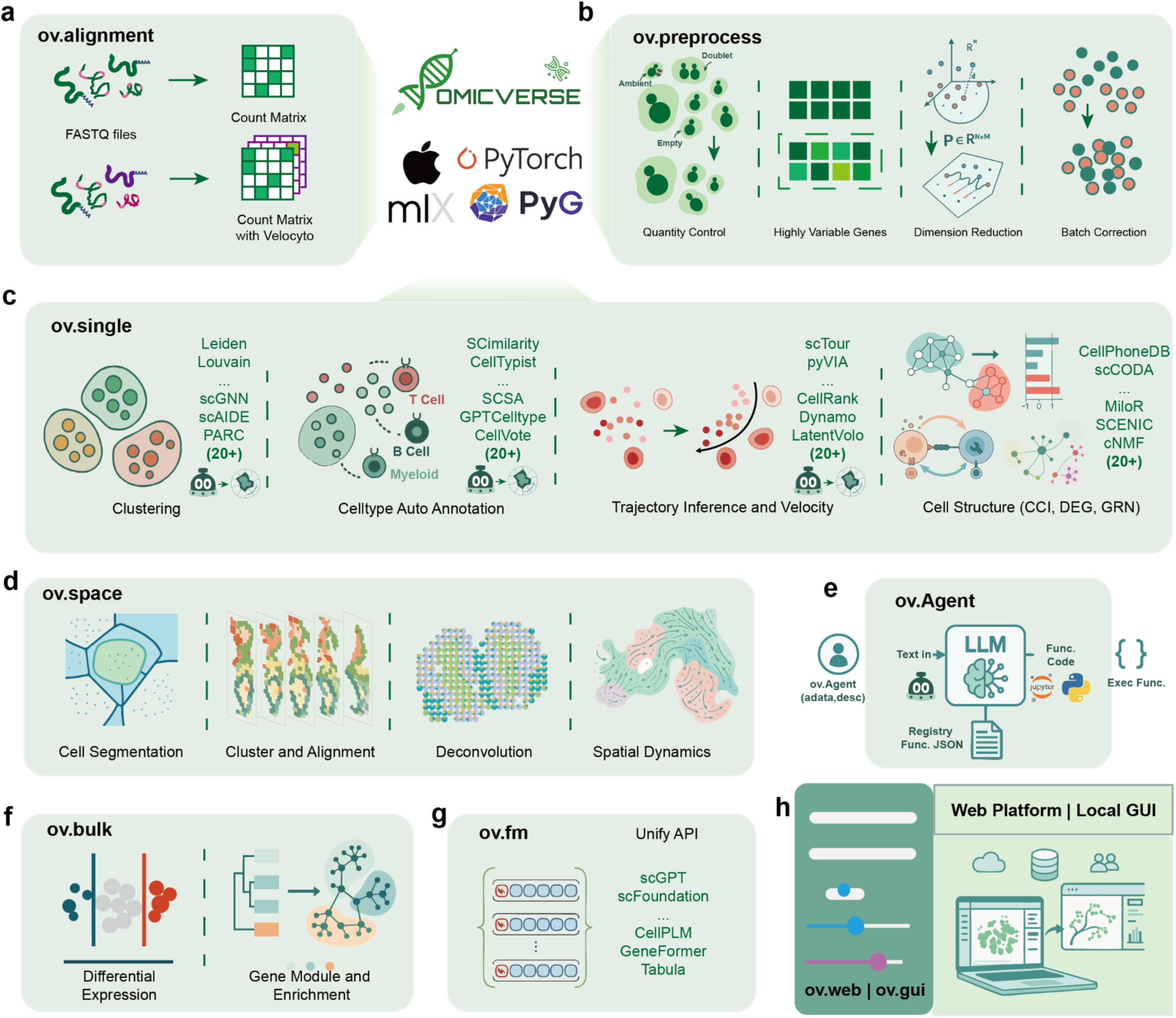
OmicVerse provides a unified ecosystem for interoperable multi-omics analysis across sequencing, single-cell, spatial, bulk and agentic workflows. **a,** The ov.alignment module supports preprocessing of raw sequencing data, converting FASTQ reads into count matrices for conventional and single-cell RNA-sequencing workflows, including velocity-compatible outputs. OmicVerse is built to interface with modern computational backends and machine-learning frameworks, including PyTorch, PyG and Apple MLX. **b,** The ov.preprocess module standardizes core preprocessing steps for single-cell analysis, including quality control, highly variable gene selection, dimensionality reduction and batch correction. These functions provide a common entry point for downstream analyses while shielding users from method-specific implementation differences. **c,** The ov.single module unifies major categories of single-cell downstream analysis within a shared interface. Supported tasks include clustering, automated cell type annotation, trajectory inference and RNA velocity, and higher-order cell structure analyses such as cell–cell interaction (CCI), differential expression (DEG) and gene regulatory network (GRN) inference. Representative integrated methods are shown for each task, illustrating that OmicVerse provides access to diverse algorithms through a consistent API rather than isolated tool-specific wrappers. **d,** The ov.space module extends the same design principle to spatial omics, covering cell segmentation, cluster mapping and alignment across sections, deconvolution and spatial dynamics analysis. This enables spatial workflows to be integrated with the broader OmicVerse analytical ecosystem. **e,** The ov.Agent module provides a language-driven interface to the OmicVerse ecosystem. Users supply an AnnData object together with a natural-language task description, which is parsed by an LLM and mapped onto registered functions through a function registry. The selected functions are then executed programmatically, enabling agentic analysis over the unified OmicVerse function space. **f,** The ov.bulk module supports bulk transcriptomics analysis, including differential expression testing as well as gene module discovery and functional enrichment analysis. **g,** The ov.fm module provides a unified API for large biological foundation models and language models, including representative models such as scGPT, scFoundation, CellPLM, GeneFormer and Tabula, allowing these models to be incorporated into analysis workflows through the same interoperable framework. **h,** OmicVerse is further deployed through user-facing interfaces including ov.web and ov.gui, which provide web-based and local graphical environments for interactive analysis, visualization and result exploration

OmicVerse integrates with broader Python scientific computing ecosystem and is designed to support both classical bioinformatics workflows and modern accelerated computation. Its structured, task-oriented API follows a task-oriented design that enables users to work directly with widely adopted data abstractions, including AnnData objects, NumPy arrays, sparse matrices and tabular metadata, thereby minimizing costly conversions across workflows. OmicVerse is further interoperable with machine-learning ecosystems including PyTorch, PyTorch Geometric and MLX, allowing users to connect omics analysis with scalable modeling, graph learning and foundation-model workflows (**Supplementary Figure 1**). To improve accessibility for beginners and users without coding experience, we further developed a web platform built entirely on the OmicVerse algorithmic ecosystem, with optimized visualization support for million-scale single-cell datasets (**Supplementary Figure 2-5**).

Since its initial release, OmicVerse has supported a growing body of studies, workflows and downstream tool developments across bulk, single-cell and spatial omics. Its adoption has extended beyond single-cell analysis into integrative multi-omics modeling, spatial biology, AI-assisted analytical pipelines and reproducible tutorial workflows. In the past year, two-thirds of the studies using OmicVerse were published in high-impact journals and more than 800+ stars on Github and 100k+ downloads on Pypi (Supplementary Table 3). Beyond direct scientific use, OmicVerse has also served as a foundation for reproducible notebooks, educational materials and extensible analytical pipelines, reflecting its dual role as both a research toolkit and a broader ecosystem for method development. In the context of OmicClaw, this breadth is particularly important because the value of the runtime depends on the existence of a sufficiently rich and interoperable method ecosystem underneath it.

Our team maintains OmicVerse and J.A.R.V.I.S. as a high-quality software project with strong emphasis on engineering reliability, modular design and reproducible usage. The project is supported by structured APIs, example-driven documentation, tutorial notebooks and workflow-oriented educational materials spanning different analysis modules and user entry points. These resources are designed not only to describe function usage, but also to connect analytical background, biological motivation and executable examples across bulk, single-cell, spatial and agentic workflows. Together, these practices lower the barrier to adoption for experimental researchers, reduce maintenance overhead for developers and facilitate interdisciplinary collaboration between method developers, computational biologists and end users.

## Discussion

Together, OmicVerse and J.A.R.V.I.S. define **OmicClaw** as an open and evolving platform for executable multi-omics analysis. In this architecture, OmicVerse provides the unified analytical substrate, whereas J.A.R.V.I.S. extends that substrate into a registry-grounded, state-aware and recoverable execution environment that can be accessed through Python, agentic interfaces and user-facing analytical workspaces. This separation is conceptually important: the central claim of this work is not that an LLM alone can solve bioinformatics, but that a unified omics ecosystem coupled to a bounded execution runtime can make natural-language multi-step analysis substantially more executable and reproducible. As multi-omics technologies, biological foundation models and computational infrastructure continue to evolve, we anticipate that OmicClaw will serve as a durable and extensible foundation for interoperable analysis and grounded human–AI collaboration in computational biology.

## Code Availability

The omicverse ecosystem official website is http://omicverse.com/. The source code of omicverse is licensed under GPL-v3 licensed and hosted at the public GitHub repository https://github.com/Starlitnightly/omicverse. The omicverse ecosystem package is distributed free of charge via PyPI (https://pypi.org/project/omicverse/) and conda-forge channel (https://anaconda.org/conda-forge/omicverse). Comprehensive documentation of omicverse, including guidelines, API references and example usages, are available at https://omicverse.readthedocs.io/en/latest/. The OmicClaw agent system stored in https://github.com/Starlitnightly/omicclaw The JARVIS agent system tutorial could be found in https://omicverse.readthedocs.io/en/latest/Tutorials-jarvis/t_claw_cli/

## Acknowledgements

We thank a large number of community contributors for their valuable input to the omicverse project. We gratefully acknowledge Yubo Wang (Department of Atmospheric and Oceanic Sciences, University of Wisconsin–Madison) and Zhengze Yuan (School of Arts and Sciences, Rutgers University–New Brunswick) for generously providing computational resources and access to the large language model API, which were instrumental to the development of OmicVerse Agent. We also extend our sincere thanks to Shih-Ying Yeh (Department of Computer Science, National Tsing Hua University; Comfy Org) for graciously providing dedicated server infrastructure, ongoing maintenance support, and additional access to the large language model API throughout this project. Their contributions were essential to enabling the iterative development, testing, and deployment of the agent system.

## Author Contributions

Z.Z. and H.D. conceived the project. Z.Z. led the overall development of the OmicVerse ecosystem and the design of J.A.R.V.I.S. X.W. led the development of the agent system, long-horizon harness design and benchmarking across different models. Z.L. led the development of the iMsg bot. Y.Z. contributed to the curation and organization of algorithms and cited literature across the OmicVerse ecosystem. C.X. and H.D. provided biological guidance for the conceptual framing and interpretation of OmicClaw. All authors contributed to software development, manuscript preparation and revision.

## Supplementary Material

## J.A.R.V.I.S design

The OV.Agent / J.A.R.V.I.S. runtime can be described as a layered control architecture that maps user requests onto bounded, provenance-aware analytical execution. Across the implementation reviewed here, the control path can be summarized as follows: user input and session context → registry-grounded capability discovery → controller-mediated multi-turn progression → runtime dispatch and code execution, with subagent orchestration, security and approval mechanisms, traceability, provenance, and workflow policy operating as auxiliary control layers. The subsections below provide a methods-oriented description of this runtime.

### 1. System entry and session context

User requests enter the runtime as natural-language instructions optionally paired with an in-memory AnnData or MuData object. The public entrypoint ov.Agent(…) instantiates OmicVerseAgent, and synchronous run() serves primarily as an adapter onto the asynchronous runtime path. The substantive execution flow proceeds through run_async(), which either executes explicit user-provided code directly or routes natural-language requests into the agentic controller loop. In this design, the user message and the active analysis object are treated as distinct but coupled inputs: the message defines intent, whereas the data object defines the current executable state of the analysis.

Session context is constructed from several stores rather than from the current prompt alone. The runtime can incorporate prior JSON session history, recent failing traces, and repository-owned workflow instructions derived from WORKFLOW.md. These sources are merged into the model-facing prompt bundle before a turn begins. The result is a session-scoped execution context that binds user intent to an evolving analysis state, a working directory, and a persistent run identity, thereby enabling iterative analysis instead of one-shot code generation.

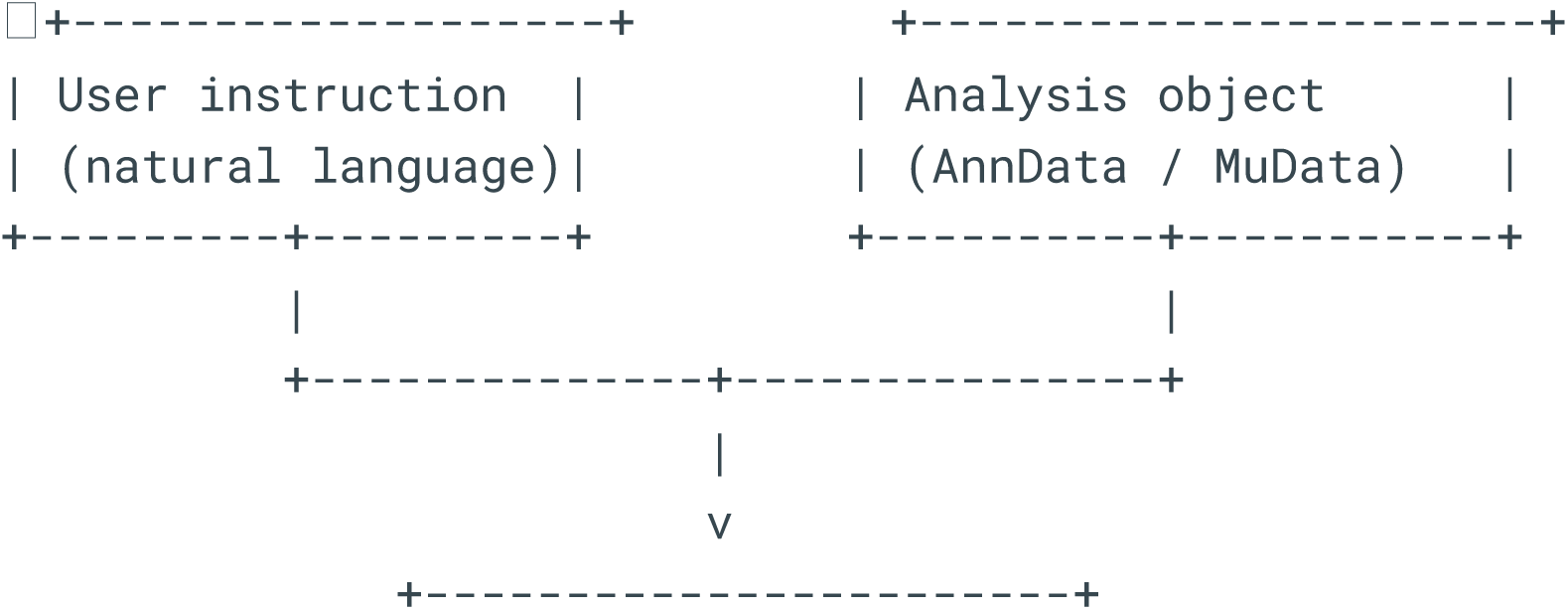
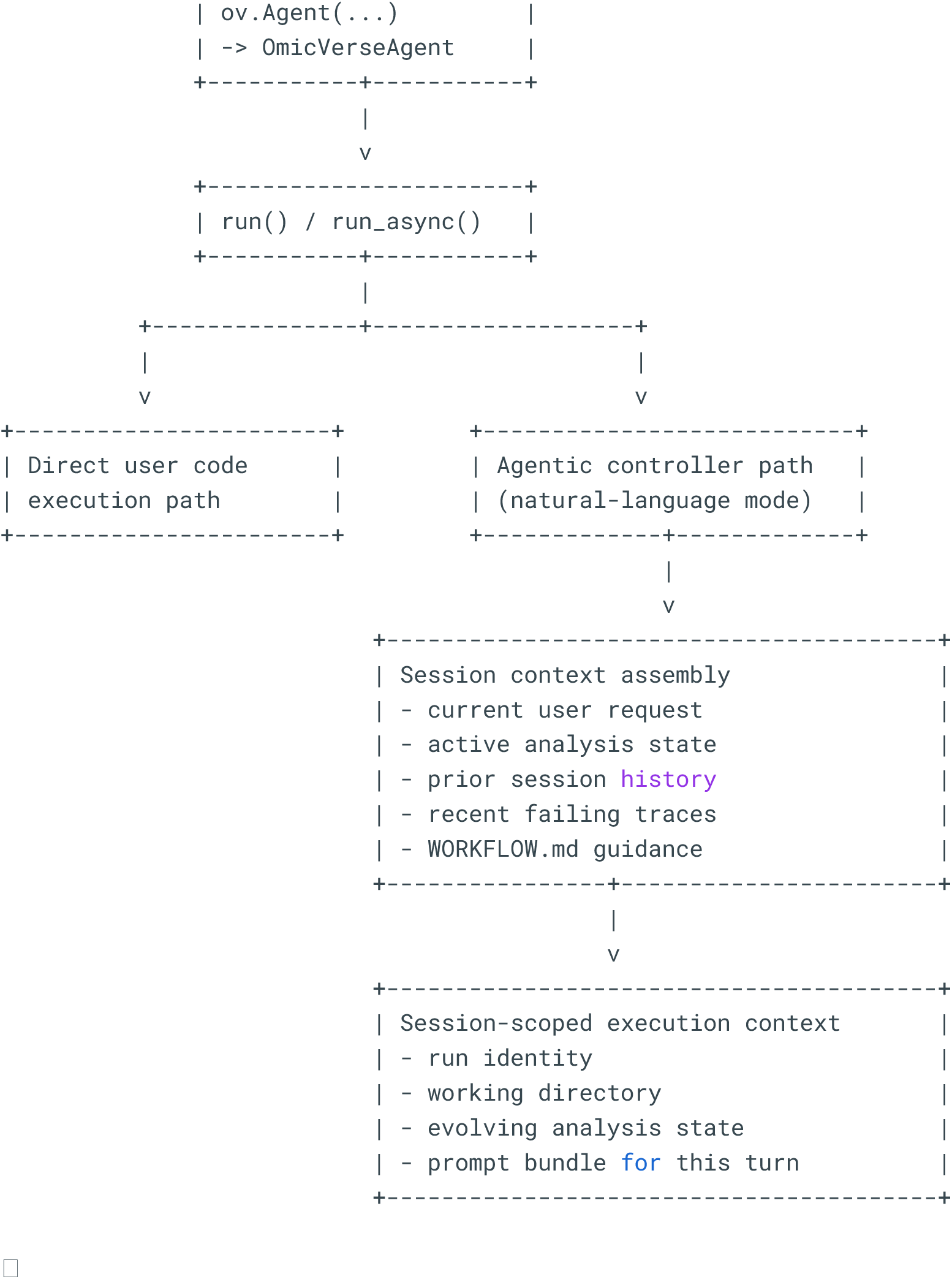

### 2. Registry-grounded capability modeling

Capability modeling is registry-grounded rather than prompt-grounded. OmicVerse functions are exposed to the runtime through @register_function and the associated function registry, which records structured metadata including canonical names, aliases, categories, natural-language descriptions, examples, prerequisite functions, required data structures, produced structures, and optional auto-fix hints. This transforms the analytical surface of OmicVerse into an inspectable action space that can be queried before execution.

The registry also supports more than exact name lookup. Retrieval logic can search aliases, descriptions, examples, imports, and derived branch-specific entries, allowing the runtime to map biologically phrased user requests onto concrete callable operations. This metadata is then used not only for search but also for weak validation: prerequisite chains can be reconstructed and required object structures can be checked against the current AnnData state. In methods terms, the registry should therefore be presented as both a discovery layer and a lightweight dependency model for executable analysis.

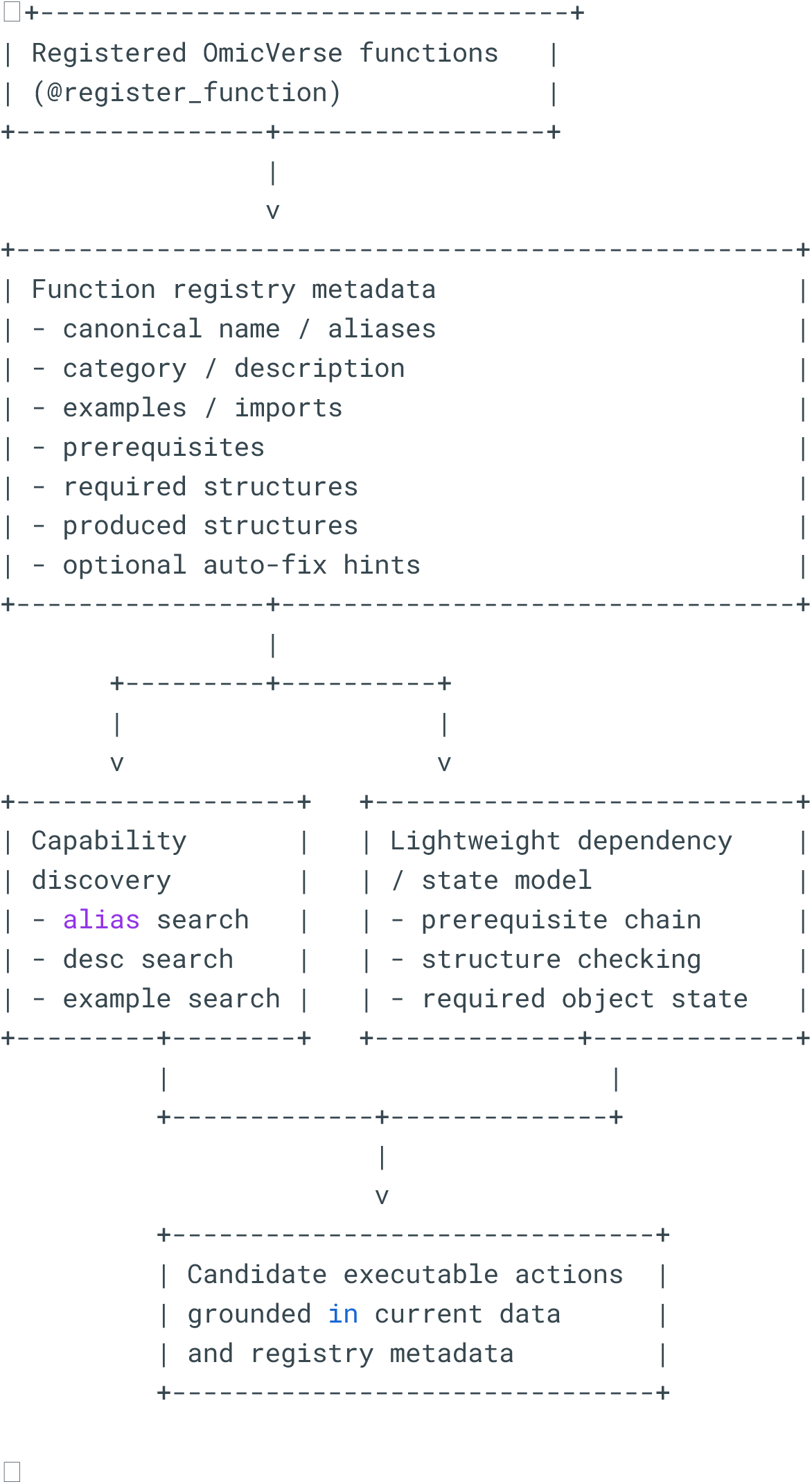

### 3. Controller-mediated multi-turn execution

Once candidate capabilities are available, the runtime advances the request through a bounded multi-turn controller loop. TurnController.run_agentic_loop() constructs the message list, exposes the currently visible tool schemas, and repeatedly invokes the model under a configurable turn budget that may be further constrained by workflow policy. Tool results are normalized back into the message stream so that subsequent turns can condition on prior actions and outcomes.

Several explicit control mechanisms regulate whether a run should continue, converge, or retry. Follow-up gating detects action-oriented requests that should not terminate in promissory text and can require the model to issue an actual tool call. Convergence monitoring detects plateaus in which the model repeatedly uses read-only tools without progressing to execution or artifact generation. In such cases, the controller can inject additional steering and continue the loop. A run terminates when the request is judged complete, when the turn budget is exhausted, or when error/cancellation conditions are reached. This makes the runtime a controller-mediated execution system rather than a free-form conversational wrapper.

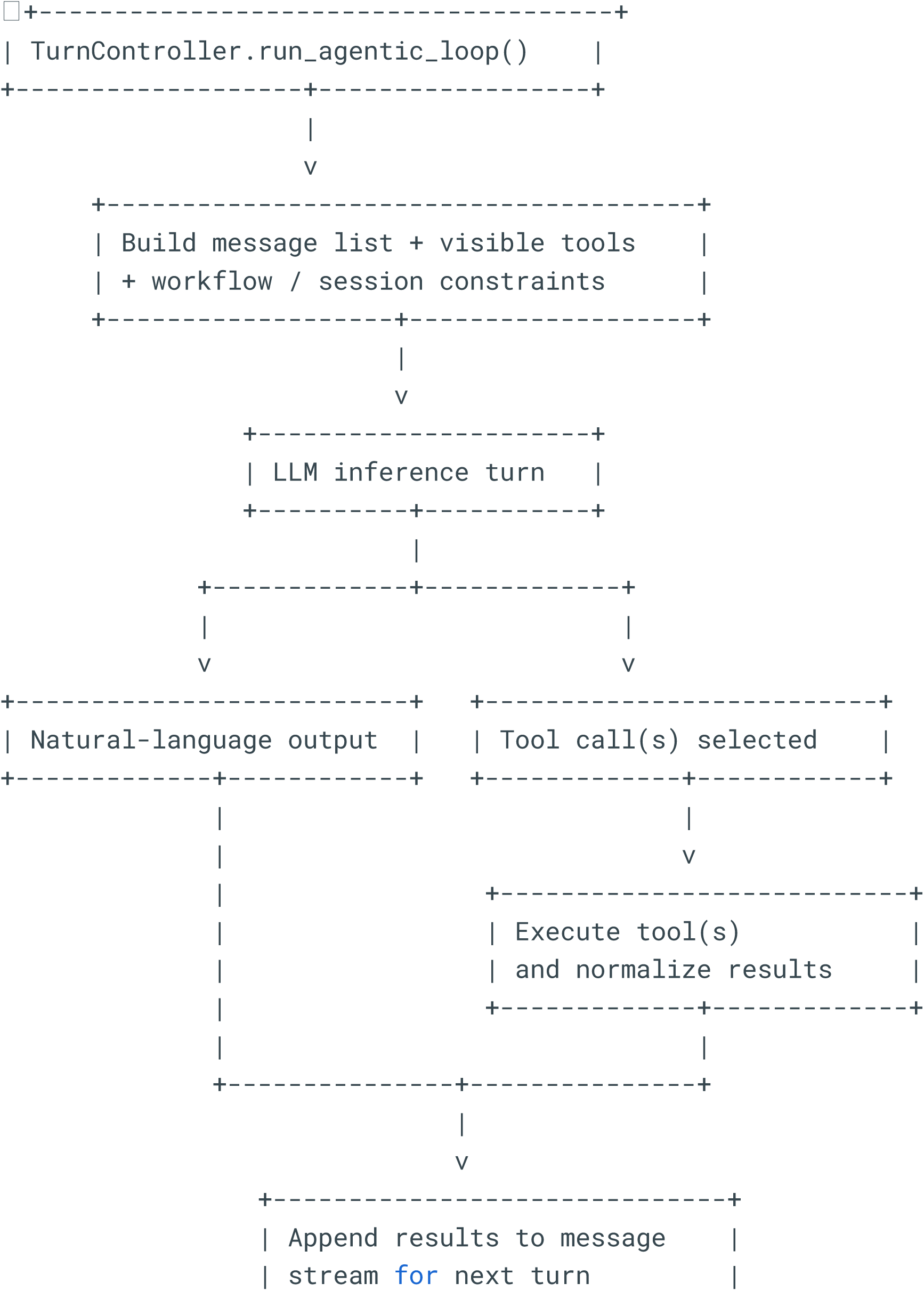
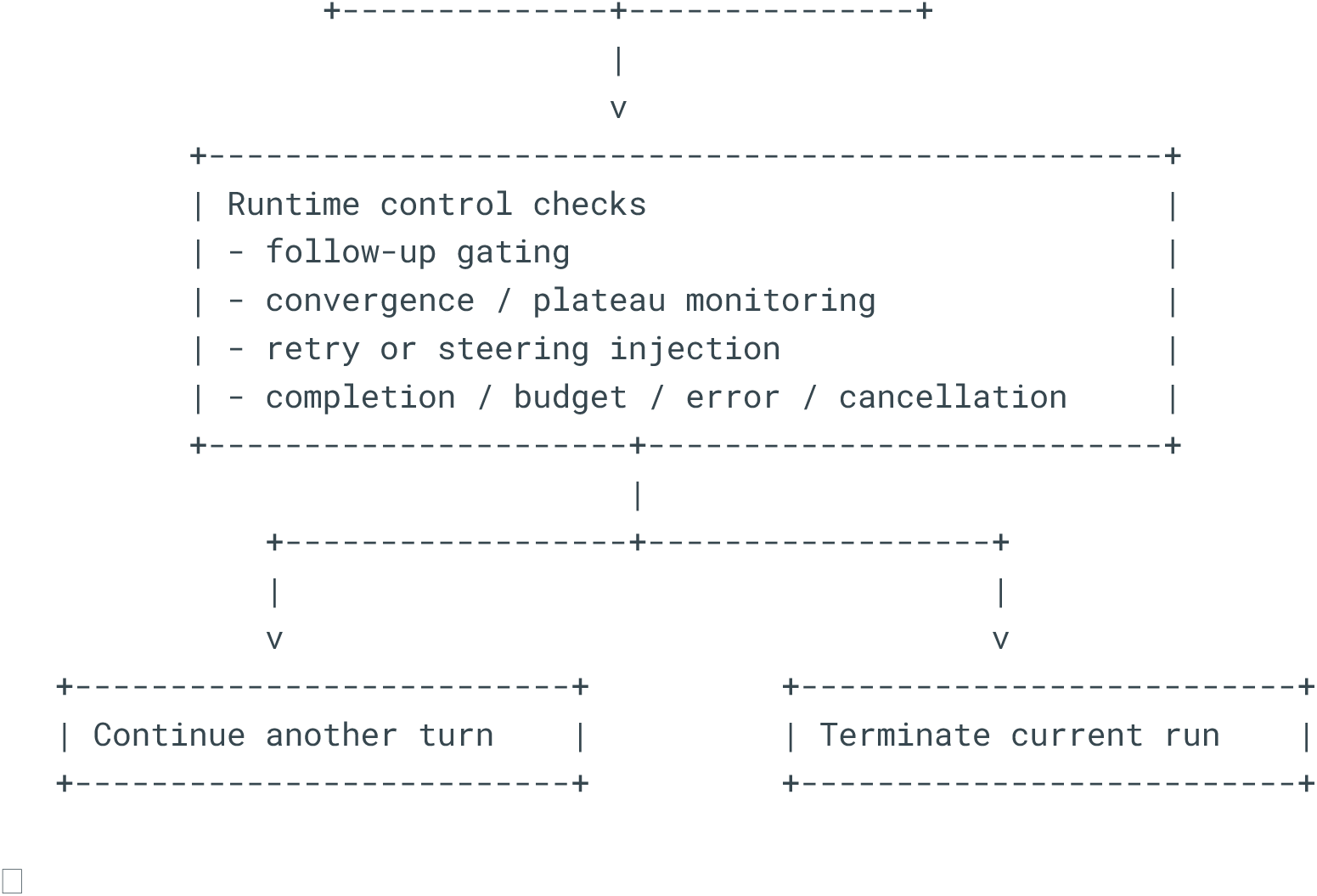

### 4. Execution runtime and code handling

Concrete operations are dispatched through ToolRuntime, which acts as the bridge between model-selected tool calls and executable handlers. This layer canonicalizes tool names, enforces runtime restrictions such as plan mode, and routes requests to OmicVerse-native inspection tools, function search, skill search, web/file operations, task and question utilities, subagent delegation, and code execution. The tool runtime therefore separates action selection from action realization.

Generated code is handled by AnalysisExecutor. Before execution, a proactive transformation stage rewrites recurrent failure patterns such as misuse of in-place OmicVerse functions or selected incompatible argument forms. The executor then performs prerequisite-aware checks using registry metadata, applies security and approval logic, and attempts execution through a notebook-backed runtime when available. If notebook execution is unavailable or fails under allowed policy, the system can fall back to restricted in-process execution using controlled globals. Error recovery is multi-stage and can include pattern-based repair, optional dependency installation, and diagnosis-guided correction. This execution path should be described as notebook-preferred with constrained fallback, rather than as unrestricted code generation.

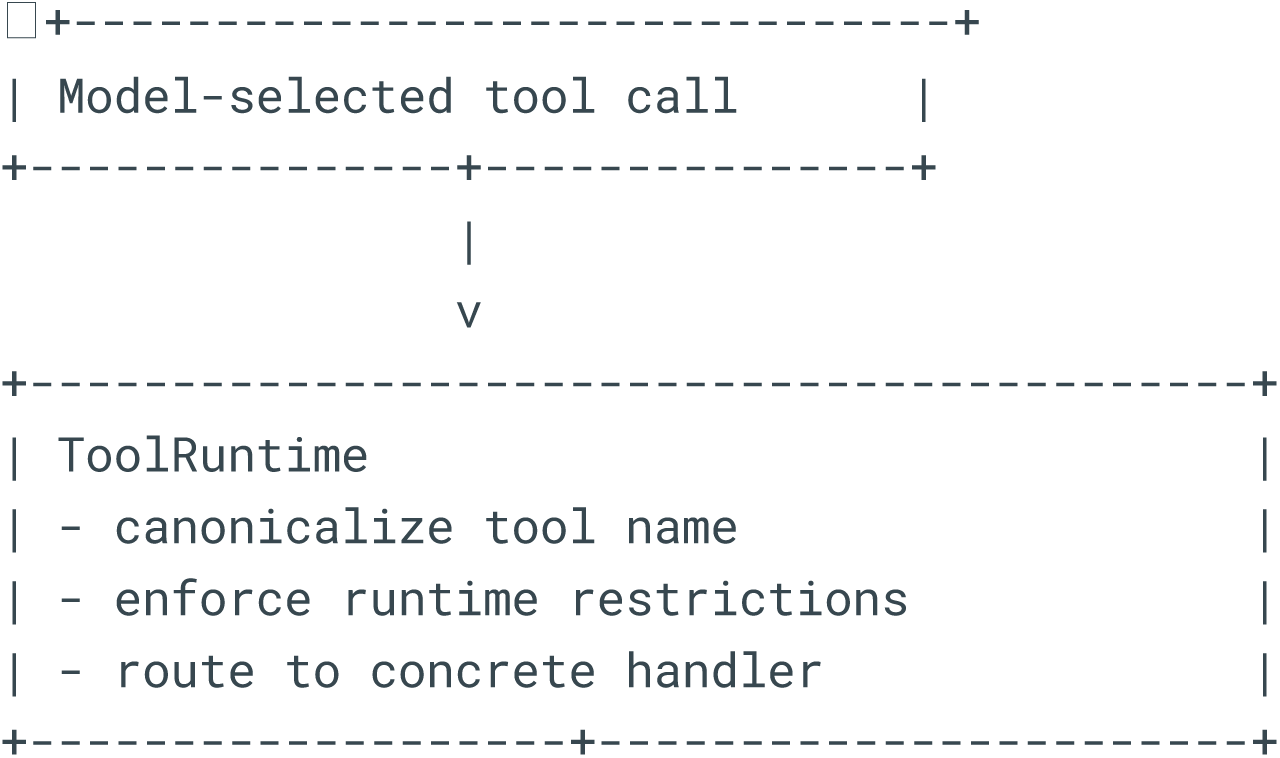
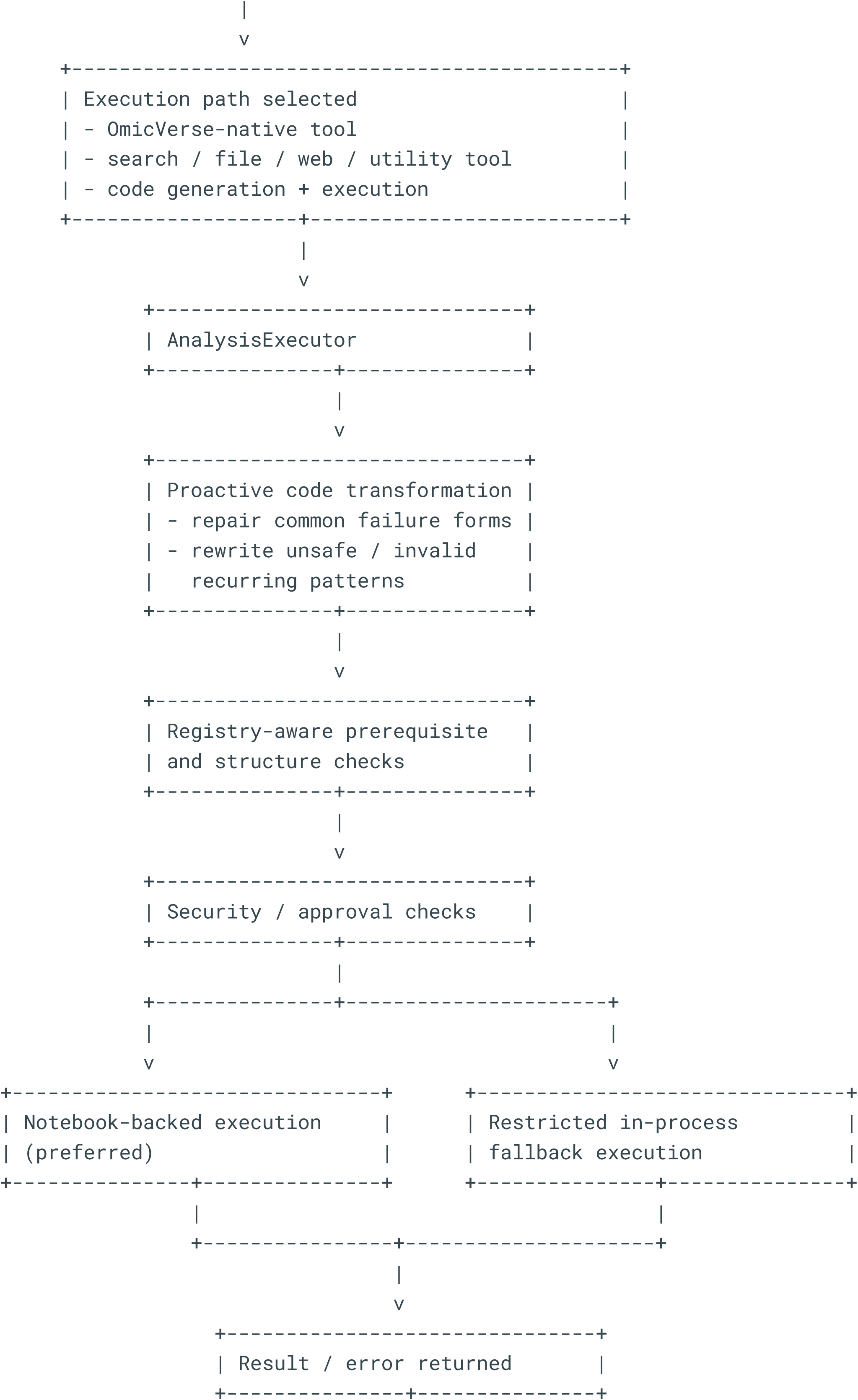
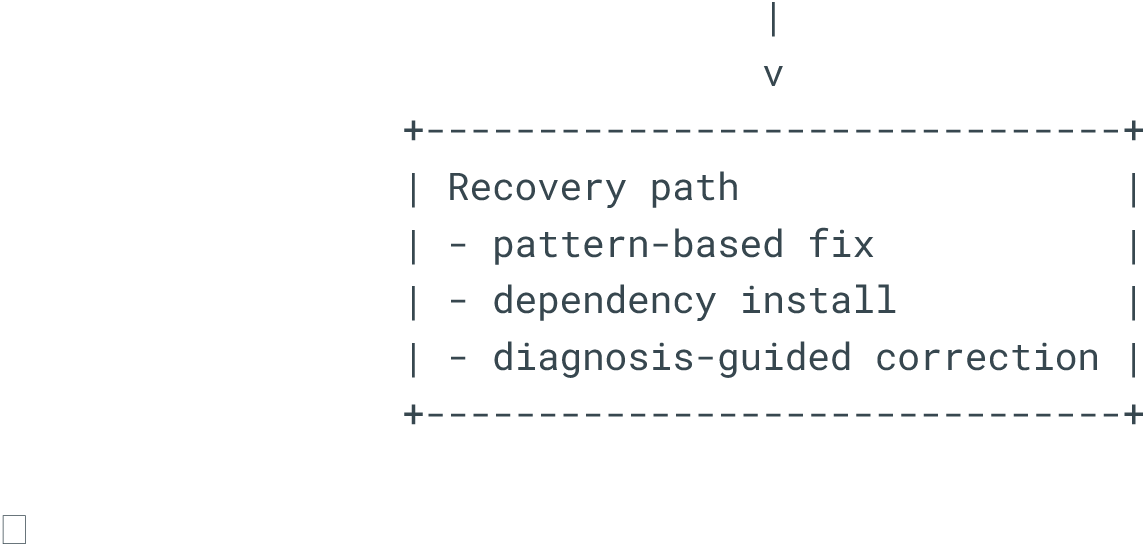

### 5. Subagent and task decomposition

The runtime supports bounded subagent delegation for task decomposition. When the parent agent invokes the subagent tool, control is transferred to SubagentController.run_subagent(), which creates a fresh prompt bundle, assigns a role-specific prompt, restricts the available tools, and runs an independent bounded loop. The built-in subagent roles separate exploration, planning, and focused execution.

Importantly, subagents do not inherit the full parent transcript as unrestricted conversational copies. Instead, they receive compact task context and operate in isolated message histories with role-limited tool surfaces. The parent runtime receives either a summary or an updated analysis object, depending on the delegated task. In methodological terms, subagents should be characterized as isolated auxiliary control loops for decomposition, not as fully independent autonomous agents.

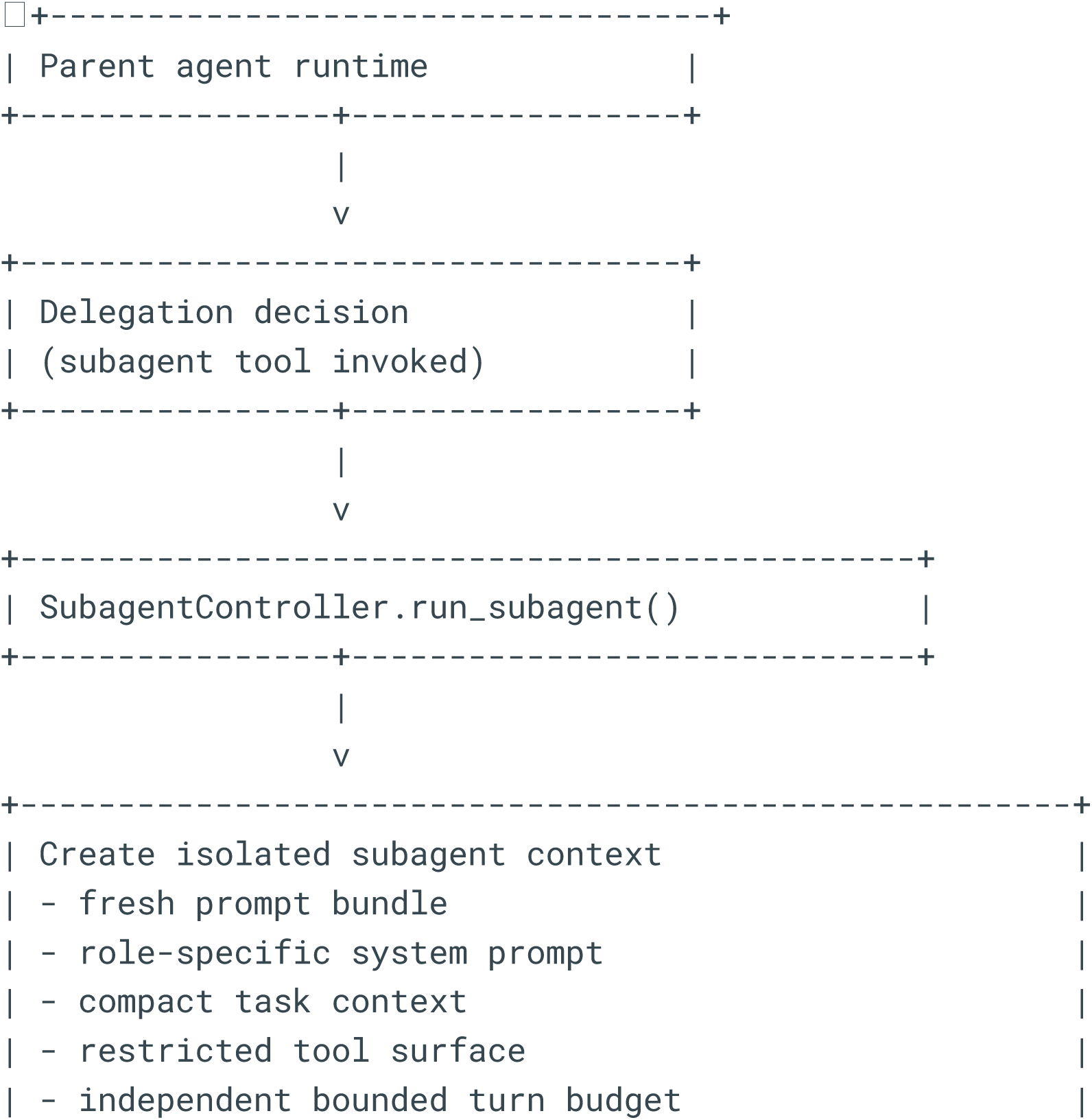
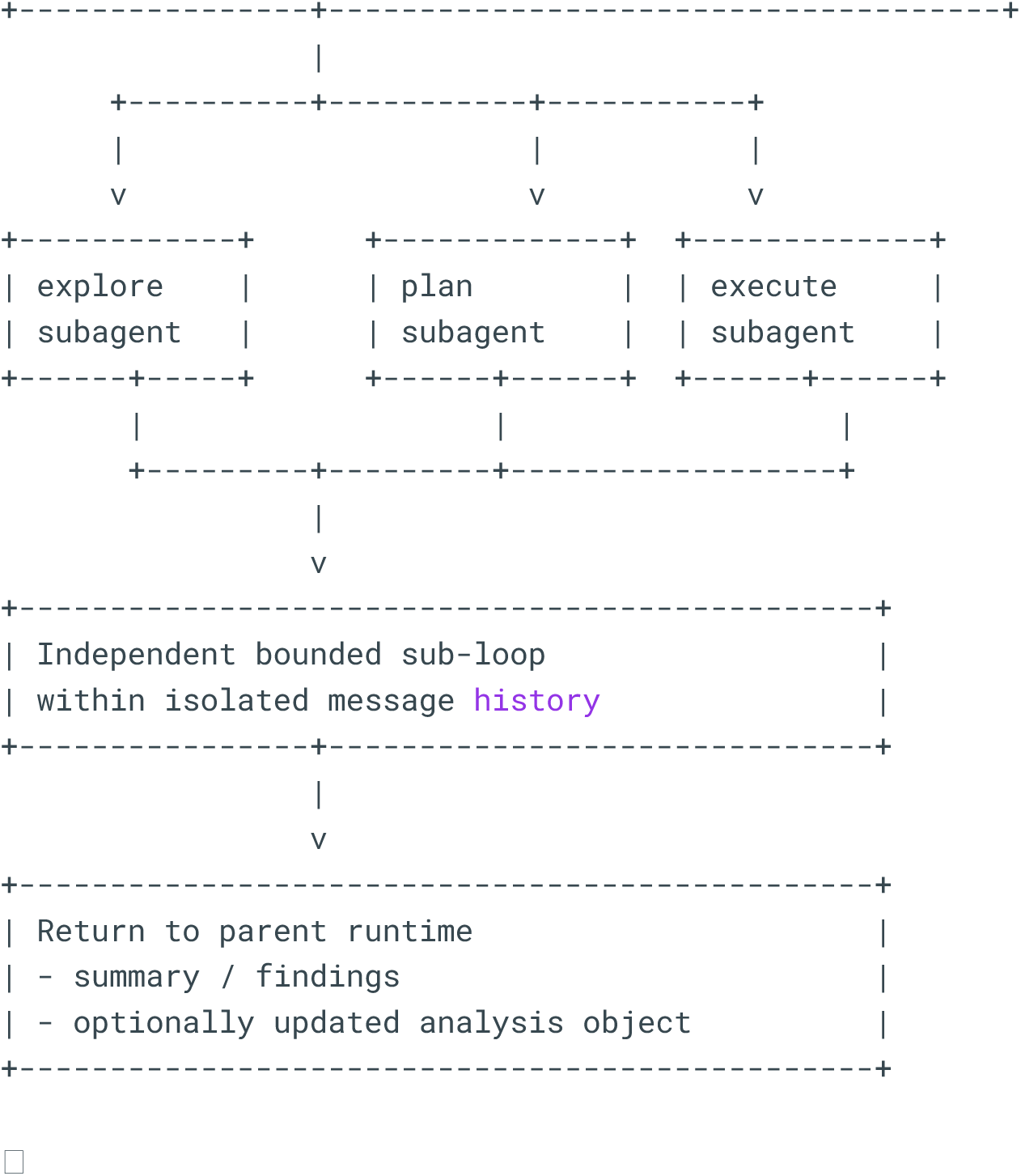

### 6. Security, policy, and approval

Security and approval are implemented as cross-cutting controls that attach to the execution path at multiple stages. Before generated code is executed, the runtime applies AST-based static scanning through CodeSecurityScanner to detect dangerous imports, blocked calls, and sandbox-escape patterns. Critical violations stop execution immediately, whereas noncritical cases may still require approval depending on ApprovalMode.

Tool-level policy is enforced separately from code scanning. High-risk tools such as shell access, file writing, notebook editing, and worktree transitions are marked as approval-gated in the tool catalog and must pass tool approval checks before dispatch. Runtime hardening is further implemented through restricted execution globals and proxies such as SafeOsProxy. The reviewed code explicitly frames this design as defense in depth rather than as a complete security boundary; the manuscript should preserve that distinction.

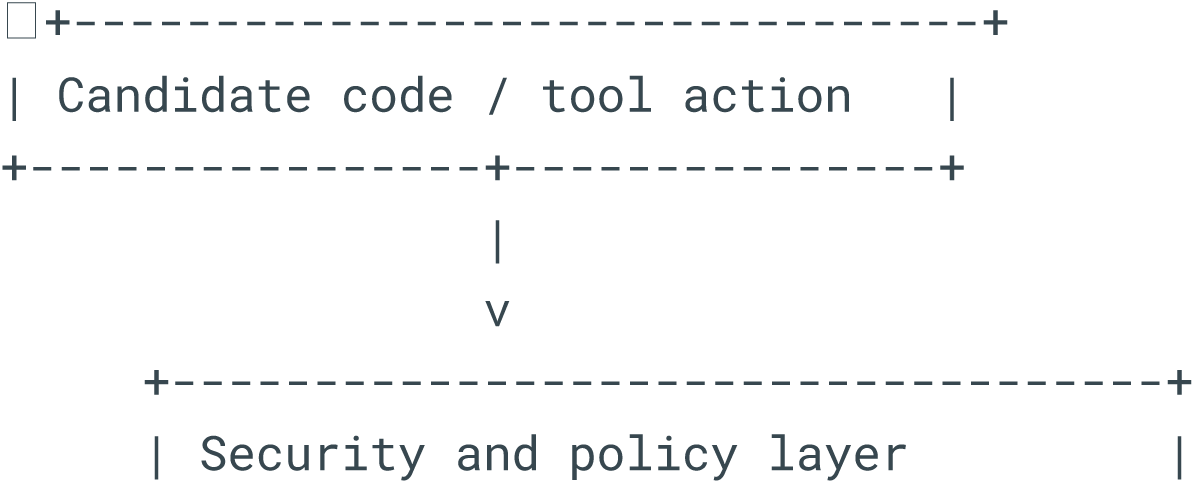
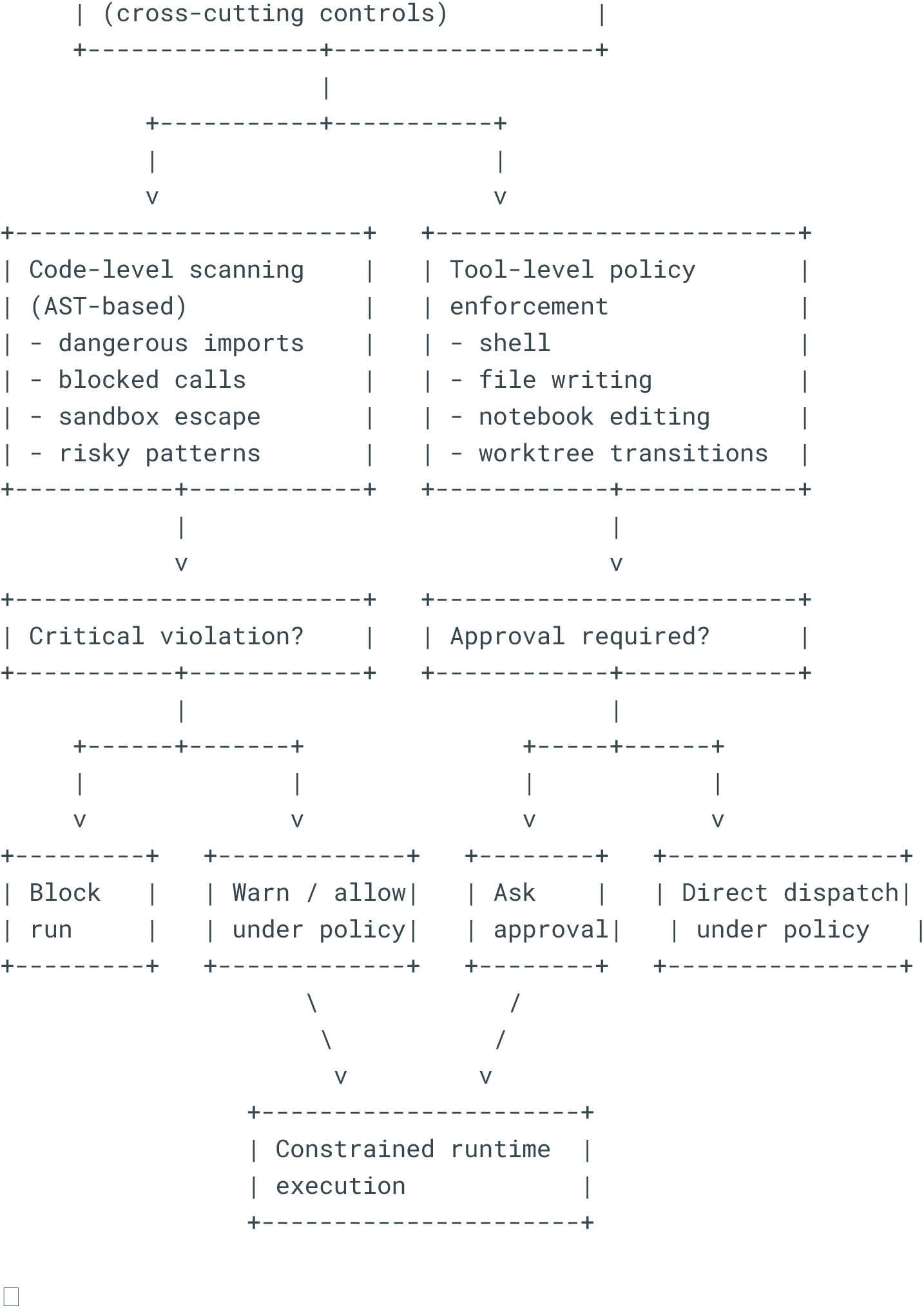

### 7. Trace, provenance, and context compaction

Provenance is maintained through structured tracing, session history, and artifact-aware run storage. During execution, RunTraceRecorder captures stepwise events, tool calls, artifacts, summaries, and final status, and RunTraceStore persists these traces on disk. After completion, the controller reconstructs a coarser session history entry that records requests, generated code, tool usage, and summarized outputs. This enables both fine-grained replay and higher-level longitudinal session memory.

Context compaction is integrated into this provenance layer rather than treated as an unrelated prompt utility. When the active prompt bundle grows too large, ContextCompactor requests a handoff-style summary that preserves function signatures, prerequisite chains, unresolved problems, and workflow constraints. The controller can then replace the larger prompt block with the compacted handoff while still retaining trace-level provenance. Artifact persistence and workflow snapshots further connect each run to figures, tables, intermediate outputs, and execution metadata, supporting post hoc inspection and reproducibility.

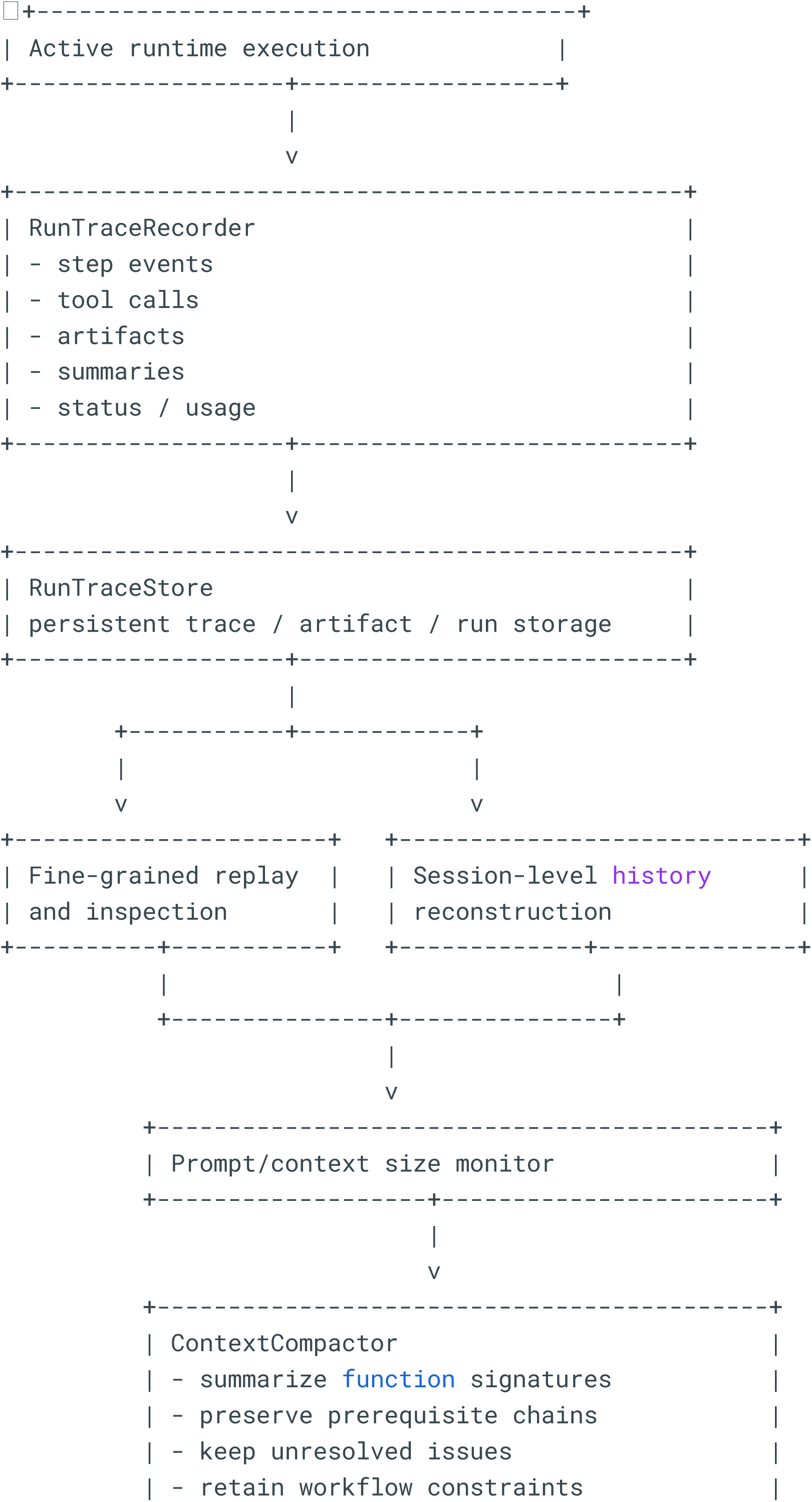
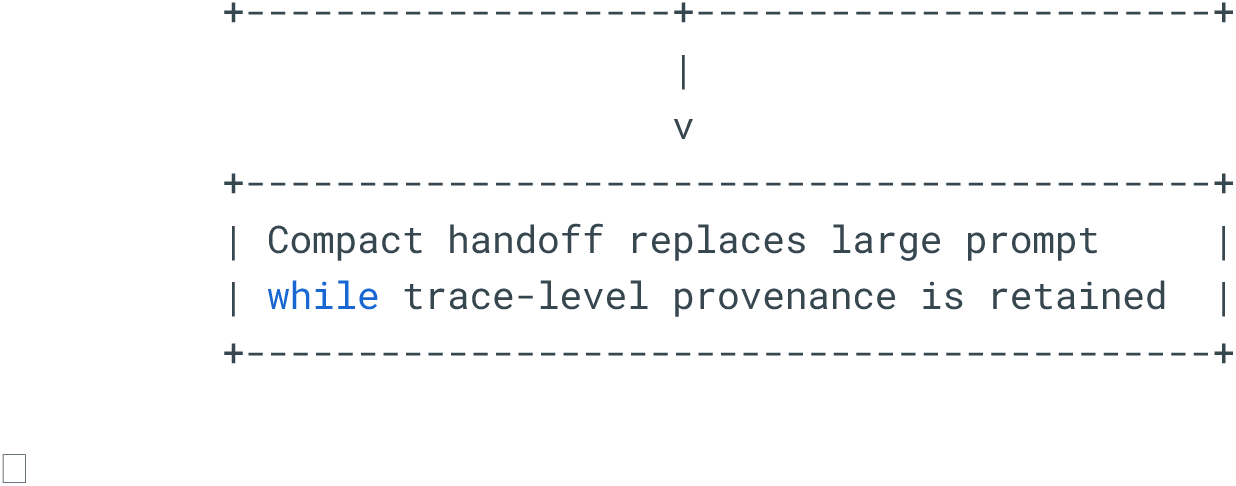

### 8. Workflow policy and extensibility

Workflow policy is repository-owned and is parsed from WORKFLOW.md into structured configuration and free-text guidance. This policy is then rendered back into prompt context and can also directly constrain runtime behavior, most clearly through loop controls such as max_turns. In parallel, run storage snapshots the workflow document into each analysis run so that the active policy is preserved together with the generated artifacts and trace linkage.

Extensibility is achieved by separating multiple control surfaces. Domain functions can be expanded through additional registry entries; workflow guidance can be extended through repository or package skill registries; and external capabilities can be surfaced through tool-catalog expansion and MCP-style exposure. This separation allows the public agent entrypoint to remain stable even as the internal analytical surface evolves. For supplementary methods, it is therefore more accurate to describe the runtime as a layered and extensible orchestration harness over OmicVerse rather than as a fixed monolithic assistant.

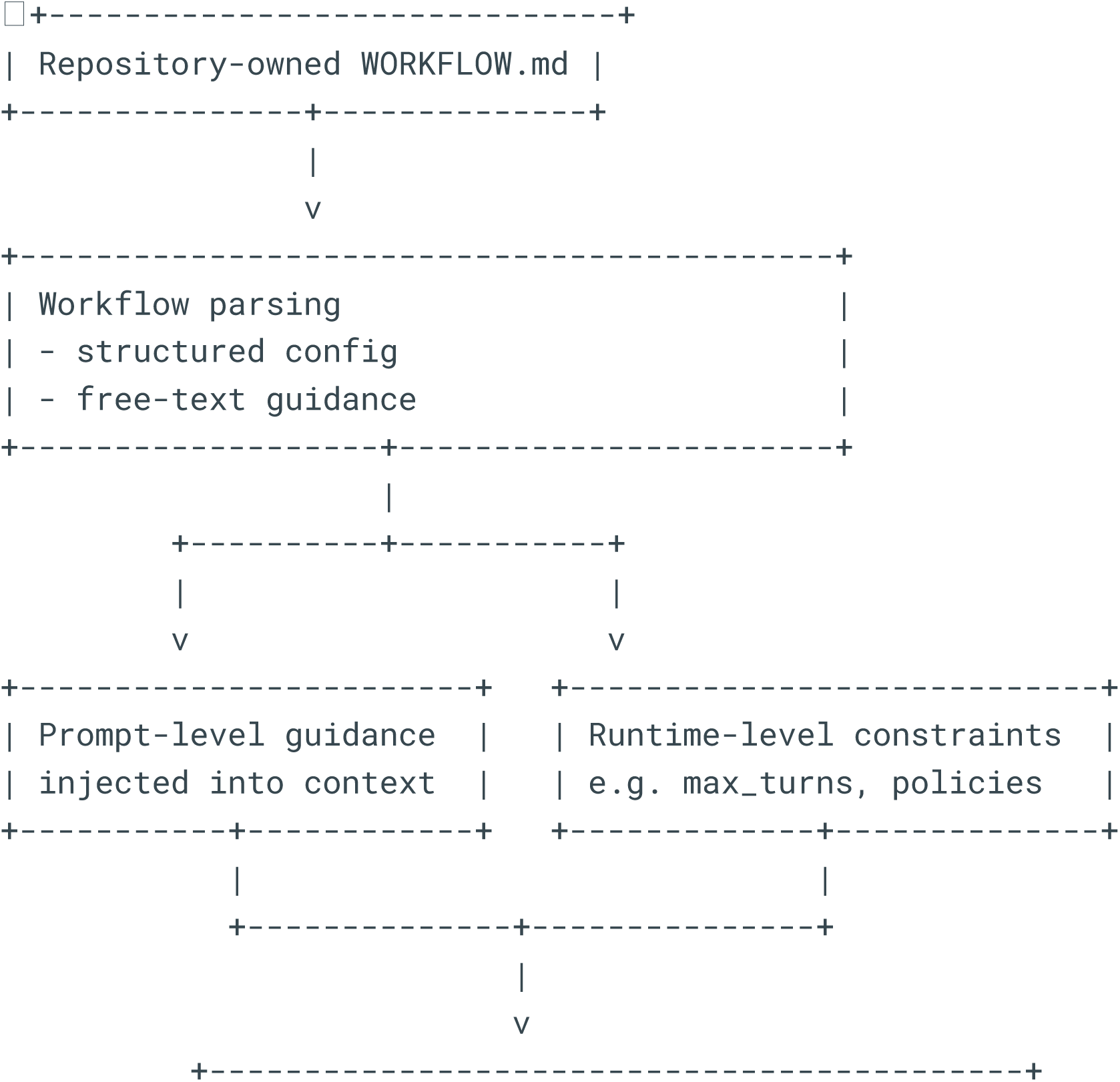
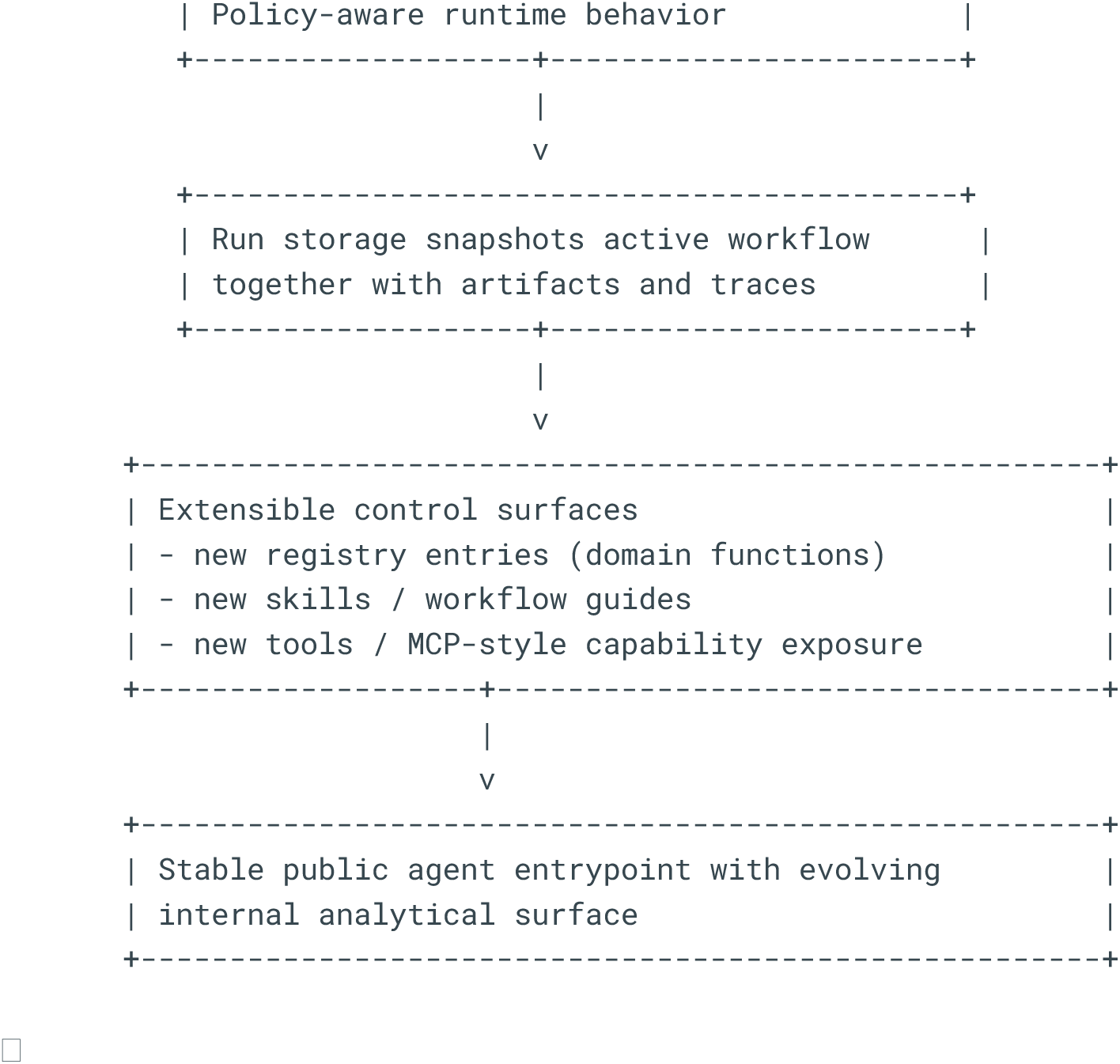

## MCP server design

The OmicVerse MCP server exposes the function registry as a set of callable tools via the Model Context Protocol (MCP), an open standard for tool-augmented AI interactions. The server communicates over standard input/output (stdio) using JSON-RPC, enabling any MCP-compatible client—including Claude Code, Claude Desktop, and custom scripts—to discover available analysis functions, inspect their parameter schemas, and invoke them without writing Python code directly. Critically, AnnData objects never cross the protocol boundary; the server holds datasets in memory and returns lightweight string handles (adata_id), so that communication overhead remains constant regardless of dataset size.

### Registry-to-manifest pipeline

At server startup, the MCP server converts the @register_function registry into a tool manifest by calling build_registry_manifest(). For each registered function, the manifest builder extracts the canonical tool name, generates a JSON Schema for its parameters, infers its execution class, and records its data requirements (prerequisite keys that must exist on the AnnData object) and return contracts (what the tool produces). The full registry contains over 200 functions and classes (Supplementary Table 2); however, only a curated subset is exposed through MCP, organized into three rollout phases. Phase P0 exposes nine core preprocessing tools (ov.io.read, ov.pp.qc, ov.pp.scale, ov.pp.pca, ov.pp.neighbors, ov.pp.umap, ov.pp.leiden, and two layer-management utilities). Phase P0.5 adds six analysis and visualization tools (marker gene detection via ov.single.find_markers and ov.single.get_markers, and four plotting functions). Phase P2 introduces five class-backed tools for differential expression (ov.bulk.pydeg), automated cell-type annotation (ov.single.pyscsa), metacell construction (ov.single.metacell), differential cell composition (ov.single.dct), and topic modeling (ov.utils.lda_topic). The phase flag (--phase) controls which tools are exposed at server startup, allowing operators to match the tool surface to the available runtime dependencies.

### Execution classes

The server classifies each tool into one of three execution classes, each of which is handled by a dedicated adapter. Stateless tools (e.g., ov.utils.read) accept file paths and return newly created AnnData handles without requiring prior session state. AnnData-backed tools (e.g., ov.pp.pca, ov.pl.embedding) accept an adata_id parameter, retrieve the corresponding in-memory AnnData object, execute the analysis function, and return the updated handle along with a state_updates record describing what was added or changed (e.g., new .obsm keys, new .obs columns). Class-backed tools (P2) implement a multi-action lifecycle: a create action instantiates the analyzer and returns an instance_id; subsequent actions (run, annotate, train, predict, or results) operate on the instance; and a destroy action frees the associated memory. This design allows stateful, multi-step analyses such as DESeq2-based differential expression to be managed within a single conversational session.

### Session model and handle management

All server-side objects are managed by a session-scoped handle store. Each session maintains three handle types: adata handles for referencing in-memory AnnData objects, artifact handles for output files on disk (plots, tables, exports), and instance handles for class-backed tool instances. Handles are isolated by session identifier, preventing cross-session access. The server enforces configurable quotas (default: 50 AnnData handles, 200 artifact handles, and 50 instance handles per session). AnnData persistence is explicit rather than automatic: the ov.persist_adata meta-tool writes the AnnData to an .h5ad file alongside a JSON sidecar containing session metadata (session identifier, creation timestamp, observation and variable column summaries), and ov.restore_adata reloads a persisted file into the current session. Instance handles are memory-only and cannot be persisted.

### Prerequisite enforcement and hallucination prevention

Each tool in the manifest declares its data requirements—the specific keys that must exist on the AnnData object before execution (e.g., ov.pp.pca requires layers[“scaled”]; ov.pp.neighbors requires obsm[“X_pca”]). Before dispatching any tool call, the executor validates these requirements against the current AnnData state. If a requirement is not met, the server returns a structured error response containing an error_code of missing_data_requirements, the specific missing keys, and a suggested_next_tools array indicating which tools should be run first. This mechanism directly prevents code hallucination: an LLM agent cannot skip preprocessing steps or invoke tools in an invalid order, because the server rejects the call before execution and provides corrective guidance.

### Observability

The server records three levels of observability data per session. Trace records capture the tool name, execution duration, input and output handle references, and success or failure status for each tool call. Event records log handle lifecycle transitions (creation, persistence, restoration, deletion) and quota enforcement actions. Aggregated metrics provide session-level summaries including total tool calls, failure counts, and handle counts. These records enable post-hoc inspection of the full analytical provenance chain, supporting the reproducibility and auditability goals described in the main text.

### Deployment

The server supports two deployment modes over the same stdio transport. In local mode, the MCP client spawns the server as a Python subprocess (python -m omicverse.mcp), suitable for interactive analysis on a single workstation. In remote mode, the client opens an SSH connection to a remote host and launches the server process there; the JSON-RPC protocol is tunneled over SSH, while all computation and data remain on the remote machine. Both modes expose identical tool interfaces, allowing users to scale from laptop-based exploration to GPU-accelerated server-side analysis without changing the analytical workflow.

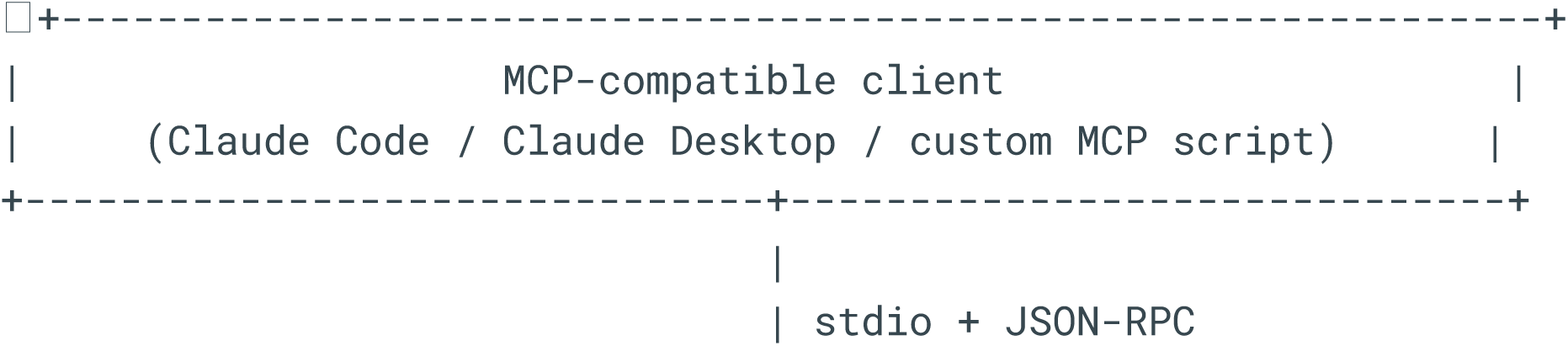
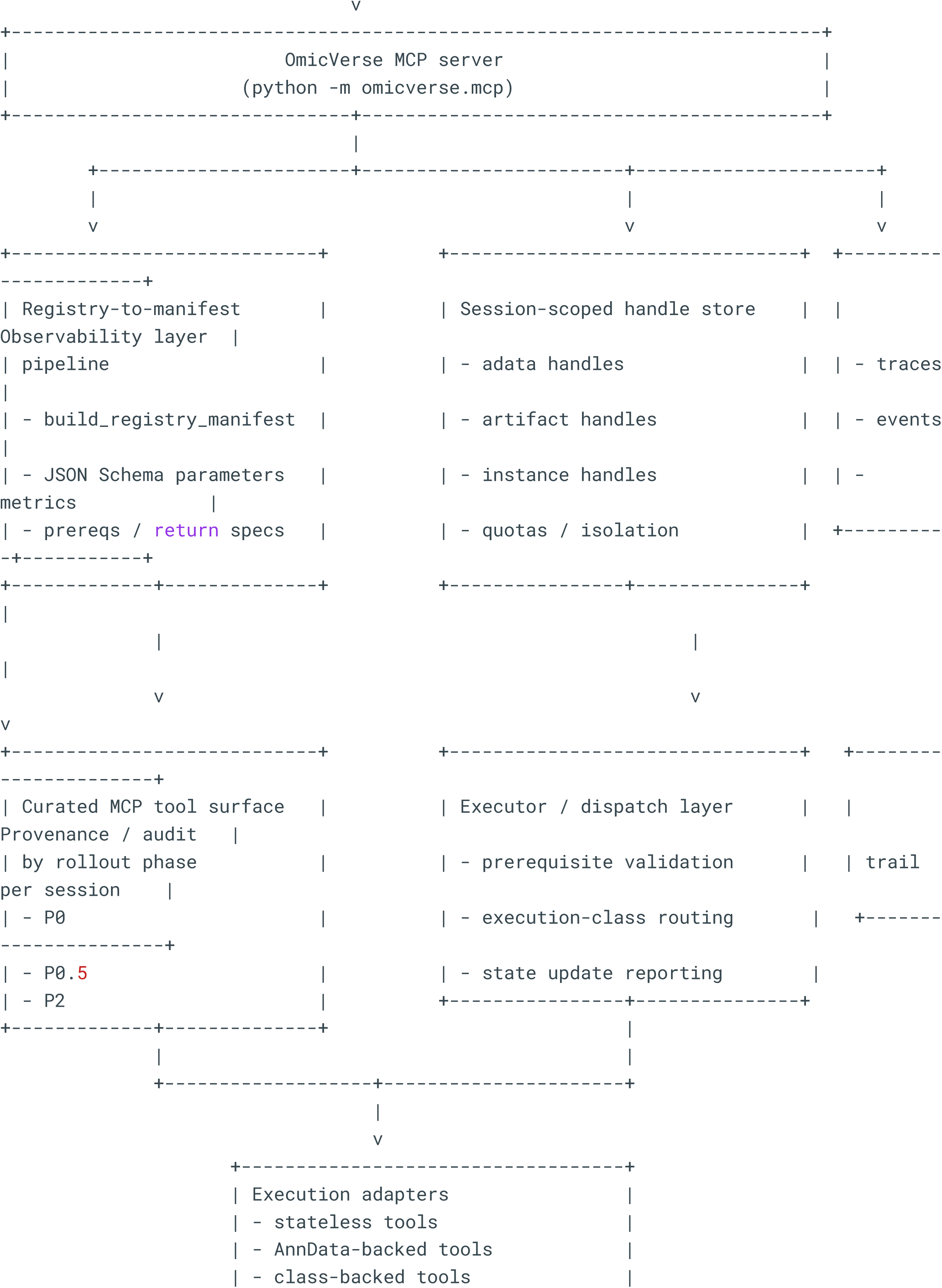
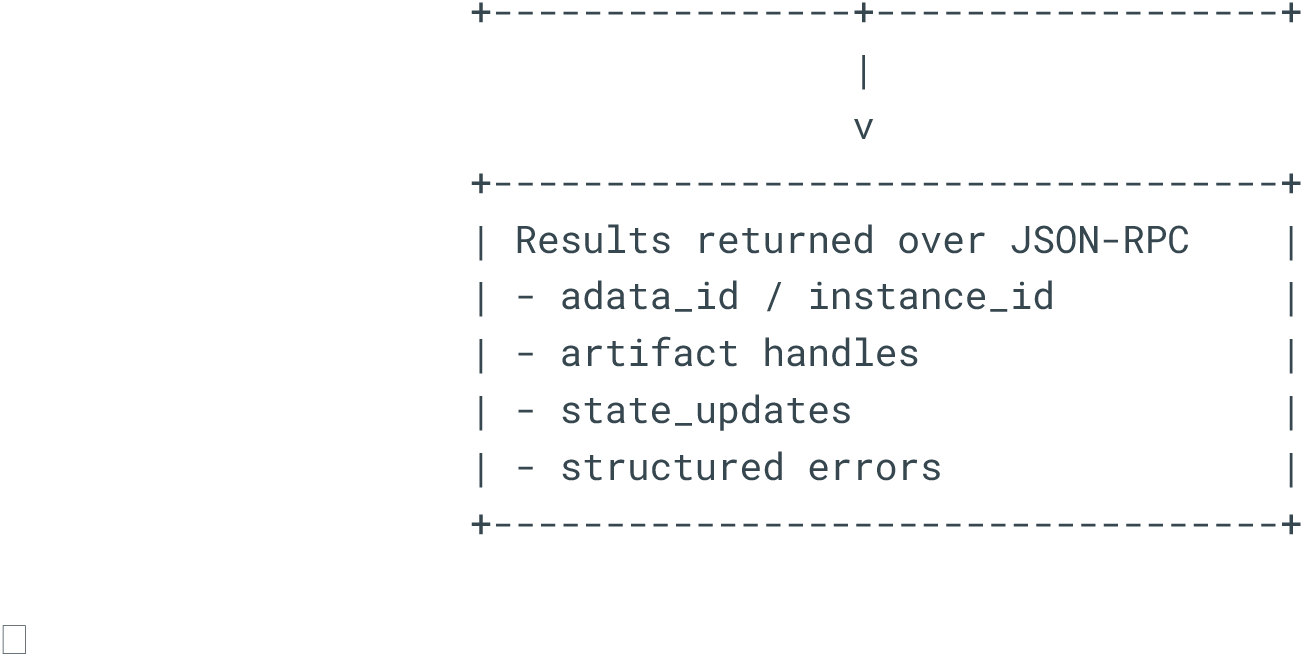

## Benchmark design and implementation

To compare the computational performance and output fidelity of the OmicVerse and Scanpy pipelines, we performed a stepwise benchmark across increasing dataset sizes using a common AnnData-based workflow. All analyses were run with a fixed random seed (RANDOM_SEED = 0) to minimize stochastic variation. Subsampled datasets contained 1,000, 2,000, 5,000, 10,000, 20,000, 50,000, and 100,000 cells, although the 1,000-cell condition was omitted from the final visualization panels for clarity. For each cell-count condition, the same input cells were processed independently by three execution modes: **Scanpy**, **OmicVerse CPU**, and **OmicVerse CPU-GPU-mixed**. The latter denotes the hybrid backend evaluated in the benchmark and was treated as a separate execution condition throughout.

The pipeline consisted of the standard single-cell preprocessing and manifold-learning sequence: library-size normalization, log transformation, highly variable gene (HVG) selection, feature scaling, principal component analysis (PCA), k-nearest neighbor (kNN) graph construction, UMAP embedding, and Leiden clustering. Benchmark parameters were held constant across methods, including selection of the top 2,000 HVGs (HVG_TOP = 2000), use of 50 principal components (N_PCS = 50), and construction of a 15-nearest-neighbor graph (N_NEIGHBORS = 15). For scaling, ZERO_CENTER = False was used. Step-level runtime was recorded in seconds for each method and dataset size and stored in a structured table (speed_df). Final processed AnnData objects were also saved for downstream metric evaluation (save_df).

Because the notebook uses a fixed seed and does not average repeated runs, each plotted point corresponds to a single deterministic benchmark execution rather than a mean across replicates.

### Runtime analysis

Runtime benchmarking was performed separately for each pipeline step. For a given method *m*, step *s*, and dataset size *n*, we denote the wall-clock time as

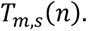

The main figure emphasizes the five computationally informative steps-HVG selection, PCA, neighbor graph construction, UMAP, and Leiden clustering-while additional preprocessing steps were retained in the full benchmark tables.

To quantify growth with dataset size, runtime scaling was modeled by a log-log linear fit,

*log*_10_ *T_m,s_*(*n*) = *α_m,s_ log*_10_ *n* + *β_m,s_*, *α_m,s_* is the empirical scaling exponent. An exponent near 1 indicates approximately linear growth, whereas larger values indicate super-linear scaling. In the displayed figure, runtime-versus-cell-count curves are shown on a logarithmic x axis.

### Reference-based evaluation of HVG and PCA outputs

For steps whose outputs can be compared directly to a reference solution, Scanpy was used as the reference implementation. Thus, OmicVerse CPU and OmicVerse mixed were evaluated against the Scanpy result for the same cell subset.

#### HVG concordance

Let *G_ref_* denote the set of HVGs selected by Scanpy and *G_ref_* the set selected by an OmicVerse backend. Concordance was quantified using both the overlap count,

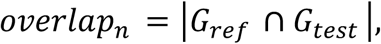

and the Jaccard index,

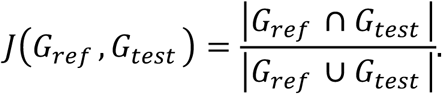

A Jaccard index of 1 indicates exact identity between the two HVG sets.

#### PCA agreement

PCA agreement was assessed by comparing the low-dimensional subspace and variance structure produced by each method to the Scanpy reference. Subspace similarity was summarized as the mean cosine of the principal angles between the reference and test PCA subspaces. If *U_ref_* and *U_test_* are orthonormal basis matrices spanning the first *k* principal-component subspaces, and *θ*_*i*_ are the principal angles between them, then the reported subspace similarity can be written as

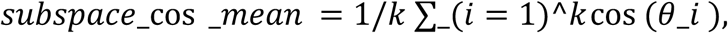

with values near 1 indicating nearly identical PCA subspaces.

Variance-ratio agreement was quantified by the mean absolute error between the explained variance ratio vectors of the two PCA results:

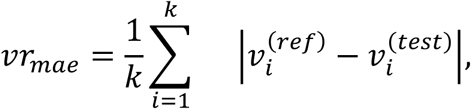

where *v_i_* is the explained variance ratio of principal component *i*.

### Intrinsic evaluation of neighborhood graphs and UMAP embeddings

For graph and embedding quality, metrics were calculated directly from each saved AnnData object rather than only against Scanpy outputs. The evaluation function used the first 50 PCs ( X_pca ) as the high-dimensional reference representation when available; otherwise, it fell back to adata. X . UMAP coordinates were taken from adata.obsm [“X_umap”]. For efficiency, up to 5,000 cells were randomly sampled per dataset for local-structure evaluation, and up to 30,000 random cell pairs were sampled for distance-based analyses.

#### High-dimensional neighborhood preservation

For each sampled query cell *i*, three neighborhoods were defined: the exact Euclidean kNN set in PCA space *H_i_*, the graph-derived neighbor set *G_i_* extracted from adata.obsp [“distances”], and the Euclidean kNN set in UMAP space *U_i_*. Preservation of local structure in the graph was summarized by the Jaccard overlap between *H_i_* and *G_i_*,

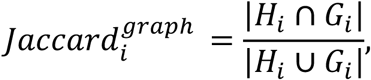

as well as by the cosine similarity between each cell and its graph neighbors in PCA space. After ℓ_2_-normalization of the high-dimensional feature vectors, this was computed as

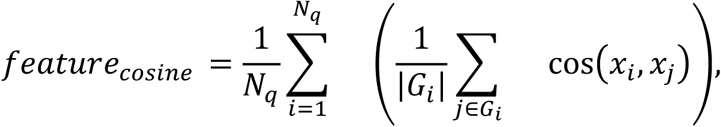

where **N*_q_* is the number of sampled query cells and *x_i_*_*i*_is the PCA-space representation of cell *i*. Higher values indicate that graph edges connect cells that remain similar in the original feature space.

Graph connectivity was further summarized by the fraction of isolated cells,

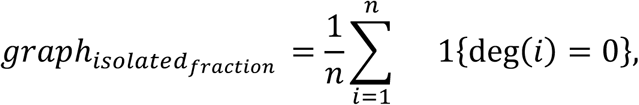

where deg(*i*) is the graph degree of cell *i*. A value of 0 indicates that every cell participates in the graph.

#### UMAP faithfulness to high-dimensional structure

UMAP quality was assessed in two complementary ways. First, local neighborhood preservation was quantified by the overlap between the high-dimensional kNN set *H_i_* and the UMAP-space kNN set *U_i_* :

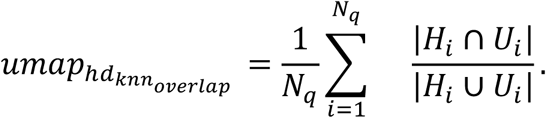

Second, global distance monotonicity was measured by the Spearman correlation between sampled pairwise Euclidean distances in PCA space and corresponding distances in UMAP space:

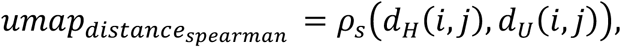

where *d_H_*(*i*, *j*) and *d_ij_*(*i*, *j*) are distances between cells *i* and *j* in PCA space and UMAP space, respectively, and *p_s_* denotes Spearman’s rank correlation. Higher values indicate that relative pairwise geometry is better preserved after embedding.

### Leiden clustering evaluation

Leiden clustering performance was evaluated from the final saved AnnData objects. Ground-truth cell annotations were taken from adata.obs [“celltype_update”], which was present in the benchmarked dataset and used as the target label set for all methods. For each method and cell-count condition, the benchmark intersected shared cell barcodes across outputs and compared predicted Leiden clusters to the groundtruth labels.

The notebook recalculated several clustering agreement metrics, of which adjusted mutual information (AMI) and homogeneity are shown in the main figure. If *C* denotes the true labels and *K* the predicted Leiden clusters, then homogeneity is defined as

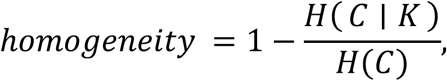

which reaches 1 when each predicted cluster contains only one true class. AMI summarizes mutual information after correcting for agreement expected by chance:

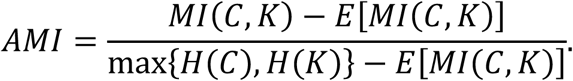

The benchmark also computed ARI, NMI, V-measure, completeness, Fowlkes-Mallows index, purity, variation of information, and the difference in the number of clusters, but only AMI and homogeneity are displayed in the main panel for compactness.

## Benchmark design of different level LLM

To systematically evaluate the analytical utility of OmicVerse ov.Agent, we implemented a benchmark framework based on code-grounded task specifications, dataset registries, unified execution harnesses, lightweight scoring rules, and reproducibility-oriented output contracts. The benchmark was designed to test whether a language-agent interface could convert natural-language task instructions into valid and traceable analysis workflows across single-cell and multimodal omics settings. Rather than evaluating isolated model responses, the benchmark assessed end-to-end task completion, including artifact generation, output validity, scientific plausibility, and, where applicable, objective agreement with ground-truth labels.

Each benchmark run was instantiated from a declarative YAML task definition containing a task identifier, dataset identifier, natural-language prompt, constraints, required outputs, scoring rubric, and time limit. During execution, the harness created a run-specific directory, loaded the requested dataset as an in-memory AnnData or MuData object, rendered the task prompt together with an explicit output contract, invoked either the full OmicVerse agent or a bare large language model baseline, and then collected logs, run manifests, and structured score summaries for downstream aggregation.

### Datasets and modalities

Datasets were centrally declared in a registry and loaded through standardized dataset-specific loader functions. The benchmark included canonical scRNA-seq resources such as PBMC3k, PBMC68k-reduced, a scvi-tools PBMC reference with batch and label annotations, pancreas integration data, and Paul15, as well as modality-specific datasets for RNA velocity (dentate gyrus), spatial transcriptomics (Visium H&E crop), scATAC-seq, CITE-seq, and multiome analysis. Each dataset specification stored metadata such as modality, species, tissue, optional batch keys, optional label keys, and cache paths, enabling both controlled execution and later aggregation into tissue-level and modality-level summaries.

### Task definitions (T1–T15)

The benchmark task suite consisted of 15 tasks spanning core scRNA-seq, velocity, spatial transcriptomics, scATAC-seq, CITE-seq, and multiome workflows. Each task specified the input object type, the intended analysis objective, required output artifacts, and any objective scoring anchor when available.

#### T1 — Quality control on PBMC3k

Input: PBMC3k AnnData.

Analysis goal: Compute standard per-cell QC metrics and apply a reasonable filtering strategy for low-quality cells and genes. At minimum, the task requires total counts, detected genes per cell, and mitochondrial fraction (or an equivalent mitochondrial metric).

- Required outputs: artifacts/qc/qc_metrics.csv with columns metric and value.
- Required outputs: artifacts/qc/qc_violin.pdf.

Objective scoring point: No external ground-truth label is used. Scoring is based on successful execution, output existence/schema, and QC reasonableness checks such as ensuring filtering does not increase cell count or collapse the dataset to zero cells.

#### T2 — Preprocessing on PBMC3k

Input: PBMC3k AnnData.

Analysis goal: Perform a standard preprocessing workflow, including normalization, log-transformation, feature selection, dimensionality reduction, and neighborhood graph construction, using reproducible and explicit parameters.

- Required outputs: preprocessing artifacts defined in the task YAML, including downstream-ready processed results.

Objective scoring point: No explicit ground-truth metric is used; scoring relies on required artifact generation and correctness of produced outputs.

#### T3 — Clustering and coarse annotation on PBMC3k

Input: PBMC3k AnnData.

Analysis goal: Run preprocessing and clustering, identify marker genes per cluster, and assign coarse immune cell-type labels to clusters based on canonical markers.

- Required outputs: artifacts/annotation/markers_top.csv with columns cluster, gene, score.
- Required outputs: artifacts/annotation/cluster_celltype_map.csv with columns cluster, celltype.
- Required outputs: artifacts/annotation/umap_celltype.pdf.

Objective scoring point: No hidden ground truth is used. Evaluation is based on artifact correctness and scientific plausibility of clustering and annotation outputs.

#### T4 — Cell-type label prediction on PBMC scvi-tools dataset

Input: PBMC scvi-tools AnnData, with ground-truth label fields removed from adata.obs before benchmarking.

Analysis goal: Infer per-cell coarse PBMC labels from expression and clustering alone. Predictions must be restricted to a predefined controlled vocabulary of nine labels, including CD4 T cells, CD8 T cells, NK cells, B cells, CD14+ monocytes, FCGR3A+ monocytes, dendritic cells, megakaryocytes, and Other.

- Required outputs: artifacts/annotation/predicted_labels.csv with columns cell_id and pred_label.
- Required outputs: artifacts/annotation/umap_predicted_labels.pdf.

Objective scoring point: This task includes an explicit objective label-accuracy term. Predicted labels are matched to the hidden dataset labels through cell_id, canonicalized across common naming variants, and converted into a benchmark accuracy score.

#### T5 — Batch integration on pancreas dataset

Input: Pancreas integration benchmark AnnData.

Analysis goal: Preprocess the data, perform batch integration with an established method available in the Python ecosystem, compute a post-integration embedding, and visualize pre/post integration structure with respect to batch and, when available, cell type.

- Required outputs: artifacts/integration/integration_summary.json.
- Required outputs: artifacts/integration/adata_integrated.h5ad.
- Required outputs: artifacts/integration/umap_integration_diagnostics.pdf.

Objective scoring point: No explicit gold-standard label metric is enforced; evaluation is based on required output validity and scientific reasonableness.

#### T6 — RNA velocity on dentate gyrus

Input: Dentate gyrus velocity dataset.

Analysis goal: Run an RNA velocity workflow using available velocity-compatible fields and produce velocity-informed low-dimensional summaries.

- Required outputs: velocity-specific artifacts defined in the task contract.

Objective scoring point: No explicit external gold standard is scored; performance is evaluated by execution, artifact validity, and plausibility checks.

#### T7 — Spatial analysis on Visium H&E crop

Input: Visium spatial AnnData containing spatial coordinates and, where available, Squidpy-compatible metadata.

Analysis goal: Confirm spatial metadata, run a standard spatial workflow including normalization, PCA, neighborhood graph construction, Leiden clustering, and spatial graph computation when available, and visualize the resulting clusters in both spatial and UMAP spaces.

- Required outputs: artifacts/spatial/spatial_summary.json.
- Required outputs: artifacts/spatial/spatial_clusters.pdf.
- Required outputs: artifacts/spatial/umap_clusters.pdf.

Objective scoring point: No hidden ground truth is used; scoring is based on artifact completeness and analysis plausibility.

#### T8 — scATAC-seq analysis on PBMC 5k

Input: PBMC scATAC-seq dataset.

Analysis goal: Perform a standard scATAC workflow, typically involving TF–IDF transformation, latent semantic indexing, and downstream clustering or embedding.

- Required outputs: defined in the task-specific output contract.

Objective scoring point: No external label-based metric is required; scoring is artifact-centered.

#### T9 — CITE-seq analysis on PBMC 10k

Input: PBMC CITE-seq dataset containing RNA and protein information.

Analysis goal: Perform basic multimodal CITE-seq analysis integrating RNA and antibody-derived features and produce expected analysis artifacts.

- Required outputs: defined in the task YAML for CITE-seq benchmark outputs.

Objective scoring point: Scored through output correctness and scientific plausibility rather than a hidden-label objective.

#### T10 — totalVI on preprocessed PBMC CITE-seq

Input: Preprocessed PBMC CITE-seq MuData object.

Analysis goal: Identify RNA and protein modalities, train a lightweight totalVI model, derive a latent representation, compute UMAP and Leiden clusters in the latent space, and summarize protein markers per cluster.

- Required outputs: artifacts/totalvi/umap_totalvi.pdf.
- Required outputs: artifacts/totalvi/top_proteins_by_cluster.csv with columns cluster, protein, mean_denoised.

Objective scoring point: No explicit gold-standard task accuracy is used; scoring depends on successful multimodal modeling and valid artifact generation.

#### T11 — MultiVI on preprocessed PBMC multiome

Input: Preprocessed PBMC multiome dataset.

Analysis goal: Train or apply a lightweight joint RNA–ATAC model using MultiVI and produce a joint latent-space representation and associated outputs.

- Required outputs: defined in the task contract for multimodal modeling artifacts.

Objective scoring point: No explicit hidden-label accuracy is imposed; success is determined through contract compliance and plausibility.

#### T12 — scATAC-seq analysis on E18 mouse brain

Input: E18 mouse brain scATAC-seq dataset.

Analysis goal: Run the same core scATAC analytical logic as T8, but on a non-PBMC tissue context to test cross-tissue robustness.

- Required outputs: defined in the task YAML.

Objective scoring point: Artifact-based and plausibility-based evaluation.

#### T13 — CITE-seq analysis on CBMC 8k

Input: CBMC CITE-seq dataset.

Analysis goal: Evaluate multimodal RNA + protein handling on a non-PBMC CITE-seq dataset.

- Required outputs: defined in the task YAML.

Objective scoring point: Artifact-based evaluation without an explicit hidden-label objective.

#### T14 — Joint multiome analysis on human brain 3k

Input: Human brain multiome AnnData containing both gene-expression and peak features.

Analysis goal: Identify RNA and ATAC feature encodings, construct a joint representation either through conversion to MuData and WNN-style analysis or through a lightweight MultiVI workflow, compute UMAP and Leiden clustering, and summarize cluster sizes and pipeline parameters.

- Required outputs: artifacts/multiome_brain/mudata.h5mu.
- Required outputs: artifacts/multiome_brain/umap_joint.pdf.
- Required outputs: artifacts/multiome_brain/cluster_counts.csv with columns cluster and n_cells.
- Required outputs: artifacts/multiome_brain/pipeline_summary.json.

Objective scoring point: No explicit hidden-label metric is used; the task evaluates cross-modality interoperability and non-PBMC generalization.

#### T15 — Joint multiome analysis on E18 mouse brain 5k

Input: E18 mouse brain multiome dataset.

Analysis goal: Perform the same class of joint RNA–ATAC analysis as in T14, but in a second non-PBMC tissue and species context to assess robustness of multimodal reasoning and output generation.

- Required outputs: defined in the corresponding multiome task contract. Objective scoring point: Artifact-centered and plausibility-centered evaluation.

### Agent and bare baseline execution

The primary benchmark condition used a thin adapter around the upstream OmicVerse implementation of ov.Agent. Model identifiers and API credentials were read from configuration files or resolved from environment variables. The execution wrapper enforced task-specific wall-clock limits, captured standard output and error streams, and wrote execution logs into the run directory.

To estimate the incremental contribution of agentic orchestration, the benchmark also included a bare-model baseline. In this condition, a single LLM call received the task prompt and a fixed system prompt instructing it to generate one complete Python script. The benchmark then extracted the code block, executed it once in a prepared namespace containing the loaded dataset and scientific Python libraries, and recorded the resulting logs and artifacts. In contrast to ov.Agent, this baseline had no iterative repair loop, no tool orchestration, and no recovery from intermediate failures.

### Scoring rubric and success criteria

Scoring combined rubric-weighted components covering execution success, output correctness, scientific reasonableness, reproducibility, efficiency, and, when applicable, label accuracy. Let w1, w2, w3, w4, w5, and w6 denote the rubric weights for execution success, output correctness, scientific reasonableness, reproducibility, efficiency, and objective label accuracy, respectively. The total benchmark score for a run was computed as:

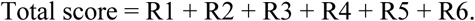

Execution success was defined as a binary term: R1 = w1 if the run completed successfully, otherwise R1 = 0. Output correctness was computed from the fraction of required artifacts that both existed and passed schema checks. If n_pass of n_required required outputs were valid, then:

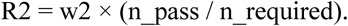

Scientific reasonableness was represented by a task-specific sanity score mapped to the R3 weight. In practice, this term was implemented through conservative heuristic checks, such as validating that QC filtering did not increase cell counts or that count-matrix outputs were not implausibly small. If s_reason denotes the raw reasonableness score normalized to [0,1], then:

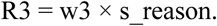

Reproducibility was awarded when the benchmark produced a valid run manifest containing the expected metadata fields. This term was treated as a fixed contribution when reproducibility metadata were successfully recorded:

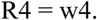

Efficiency was computed as a time-normalized fraction of the task time limit. Let t be runtime and T be the task time limit. The efficiency fraction was defined as max(0, 1 − t/T), capped to [0,1]. Thus:

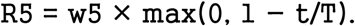

For tasks with objective labels, such as T4, per-cell accuracy was calculated by matching predicted labels to hidden ground-truth labels after canonicalization of common naming variants. If acc denotes classification accuracy, then label-accuracy points were assigned linearly as:

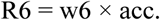

Two additional gating rules were applied. First, a run that nominally completed but failed the required artifact contract was reclassified as a benchmark failure. Second, tasks with an explicit label-accuracy component were not counted as successful if R6 = 0. These rules ensured that formally completed but scientifically empty runs did not inflate completion statistics.

### Aggregation and reproducibility

Each run generated a machine-readable manifest.json containing the task identifier, dataset identifier, seed, runtime, environment metadata, dataset fingerprint, required outputs, and detailed score breakdowns. A smaller run_result.json file was produced for downstream aggregation. Aggregation scripts scanned the canonical result tree, compiled per-run benchmark tables, and derived grouped summaries including task-level completion rate, mean total score, score dispersion, and average runtime. Additional summaries were computed across modality and tissue metadata, and an interoperability coverage table mapped successful task families onto broader capability categories.

To support manuscript traceability, the repository also defined an explicit figure–data contract linking each benchmark figure panel to its expected source-data files, generating scripts, and minimal acceptance criteria. Together with run manifests and standardized output contracts, this design enabled traceable regeneration of summary statistics and figures from benchmark outputs.

## OmicVerse Web Platform design

### 1. Visualization workspace for scalable single-cell and spatial data exploration

The visualization window in OmicVerse Web is designed as an interactive analytical workspace for large-scale single-cell and spatial transcriptomics data. On the backend, the platform uses AnnData as the core data structure and manages obs, var, obsm, layers, and gene expression matrices through a unified high-performance data adapter. Communication between the frontend and backend relies on FlatBuffers-based binary serialization rather than plain JSON transfer, thereby reducing communication overhead during large-scale data interaction. For large datasets, the system adopts a chunk-based streaming strategy, with a default chunk size of 50,000 cells for transfer and progressive rendering. It also supports a preview mode in which .h5ad files are opened in read-only mode with backed=’r’, enabling rapid inspection of structure and embeddings without fully loading the matrix into memory. To ensure responsive visualization across datasets of different sizes, the frontend implements a coordinated multi-backend rendering strategy: small to medium-sized datasets are displayed with Plotly for highly interactive scatter and statistical plots, whereas datasets exceeding a cell-count threshold are automatically switched to a deck.gl-based WebGL rendering path. When the GPU rendering path is unavailable, or when more robust large-scale static rendering is required, the platform provides Datashader or matplotlib rasterization as fallback options. Accordingly, this workspace is engineered for visualization at the million-cell scale rather than being limited to conventional small- or medium-sized single-cell data browsing. At the analytical level, the visualization workspace serves not only as a display interface but also as a complete entry point for analysis, integrating preprocessing, quality control, highly variable gene selection, dimensionality reduction, neighbor graph construction, clustering, cell-type annotation, trajectory inference, differential expression analysis, and cell–cell communication modules. This allows users to complete a closed-loop workflow of parameter setting, execution, and result inspection within a single interactive view.

### 2. Notebook-style code editor with optimized AnnData and DataFrame inspection

The code editor page adopts a notebook-style layout similar to JupyterLab, but it is not a simple replica of a traditional notebook. Instead, it is specifically optimized for the interpretability of biological data objects and for memory-aware interaction. The platform maintains a persistent Python namespace in the browser through a lightweight in-process ipykernel, allowing code cells, text files, Markdown documents, and skill files to be edited and executed within a unified session. It also supports interruption, restart, state polling, and multi-kernel context switching. To reduce frontend blocking caused by directly expanding large objects, the variable inspector implements specialized summary-display strategies for both DataFrame and AnnData objects. For DataFrames, the system returns only a limited preview of rows and columns by default, while preserving column type labels and scroll-based browsing. For AnnData, the editor presents an object summary showing the structure of obs, var, uns, obsm, and layers, and allows users to expand content slot by slot rather than serializing the complete object to the frontend at once. For matrix-like slots, only local sliced previews are returned in order to avoid densifying the entire object. The left side of the editor also includes a resource-monitoring sidebar for the active computational session, showing real-time kernel memory usage and the ranked sizes of major variables, thereby helping users identify memory-intensive objects and optimize their analysis workflow. In addition, the text editing component integrates CodeMirror-based syntax highlighting and supports common file types used in scientific workflows, including Python, R, YAML, Shell, JSON, and Markdown. For Markdown files, it provides a split-pane interface with source editing on one side and live rendered preview on the other. Together, these design choices preserve the flexibility of notebook-based analysis while substantially improving structured understanding and resource controllability for single-cell data objects.

### 3. Embedded terminal workspace for environment-level reproducibility and direct system access

The terminal page is designed as a full system workspace on the same level as the visualization and code editor interfaces, rather than as an auxiliary log window. On the backend, the platform launches a real shell process through a PTY, while the frontend continuously receives terminal output through Server-Sent Events and sends keyboard input and window-size updates through dedicated interfaces. As a result, users obtain an interaction experience in the browser that is close to a local terminal. This design enables OmicVerse Web to support not only graphical analysis but also command-line-dependent tasks such as environment management, script debugging, package installation, and system-level diagnostics, thereby covering many steps in bioinformatics workflows that cannot be meaningfully replaced by a graphical interface alone. Unlike conventional web tools that completely separate code execution from terminal access, OmicVerse Web incorporates terminal operations, kernel execution, and visualization state into a unified workspace, allowing users to switch rapidly between GUI-based analysis, free-form code execution, and shell-level operations, thereby improving the reproducibility and engineering flexibility of complex analyses.

### 4. Agent workspace for stateful, auditable, and human-in-the-loop analysis assistance

The Agent page directly embeds ov.Agent into the OmicVerse Web analytical workflow, so that agent-assisted analysis is no longer an isolated chat window but instead becomes a stateful analytical interface coupled to the current data object, code session, and file-system context. The platform uses a Server-Sent Events-based streaming mechanism to return model outputs, tool calls, code execution events, task states, and final results to the frontend in real time. At the same time, it maintains an independent session, turn, trace, and runtime state for each conversation, thereby supporting history replay, task tracking, execution cancellation, and multi-turn context retention. To strengthen controllability and auditability, the Agent workspace further implements human approval, interactive questioning, task lists, and a plan mode, allowing the model’s actions to remain bounded and reviewable when it invokes code execution, tool calls, or file operations. In this way, the Agent workspace is not merely a natural-language interaction layer, but rather a streamed, auditable, interruptible, and recoverable human–AI collaboration framework, representing one of the major engineering features that distinguishes OmicVerse Web from conventional bioinformatics web frontends.

### 5. Skill Store workspace for user-extensible method curation and workflow reuse

The Skill Store page is designed to explicitly organize reusable analytical knowledge and operational workflows in OmicVerse as locally manageable skill resources. Each skill is centered on a SKILL.md description file and may optionally include a supplementary reference.md file. The platform allows users to create, import, search, browse, and edit these skill files directly in the browser without leaving the current analytical session. When creating a new skill, the system automatically generates an initialization template and allows reference.md to remain empty, thereby lowering the barrier to building custom skills. The frontend provides both card view and list view to support content overview and batch retrieval, respectively. At the editing level, both skill and reference files can be opened directly in the platform’s built-in split-pane Markdown editor, integrating source editing and rendered preview within a single interface. More importantly, the Skill Store is not an isolated content-management page; it is aligned with the skill discovery and invocation logic of ov.Agent, enabling user-defined skills to enter actual agent workflows. By unifying method documentation, skill encapsulation, interface-level management, and executable reuse, this design makes OmicVerse Web more than just an analysis frontend: it becomes an extensible platform that supports the long-term accumulation and reuse of user analytical experience.

## Supplementary Figure

**Supplementary Figure 1.**
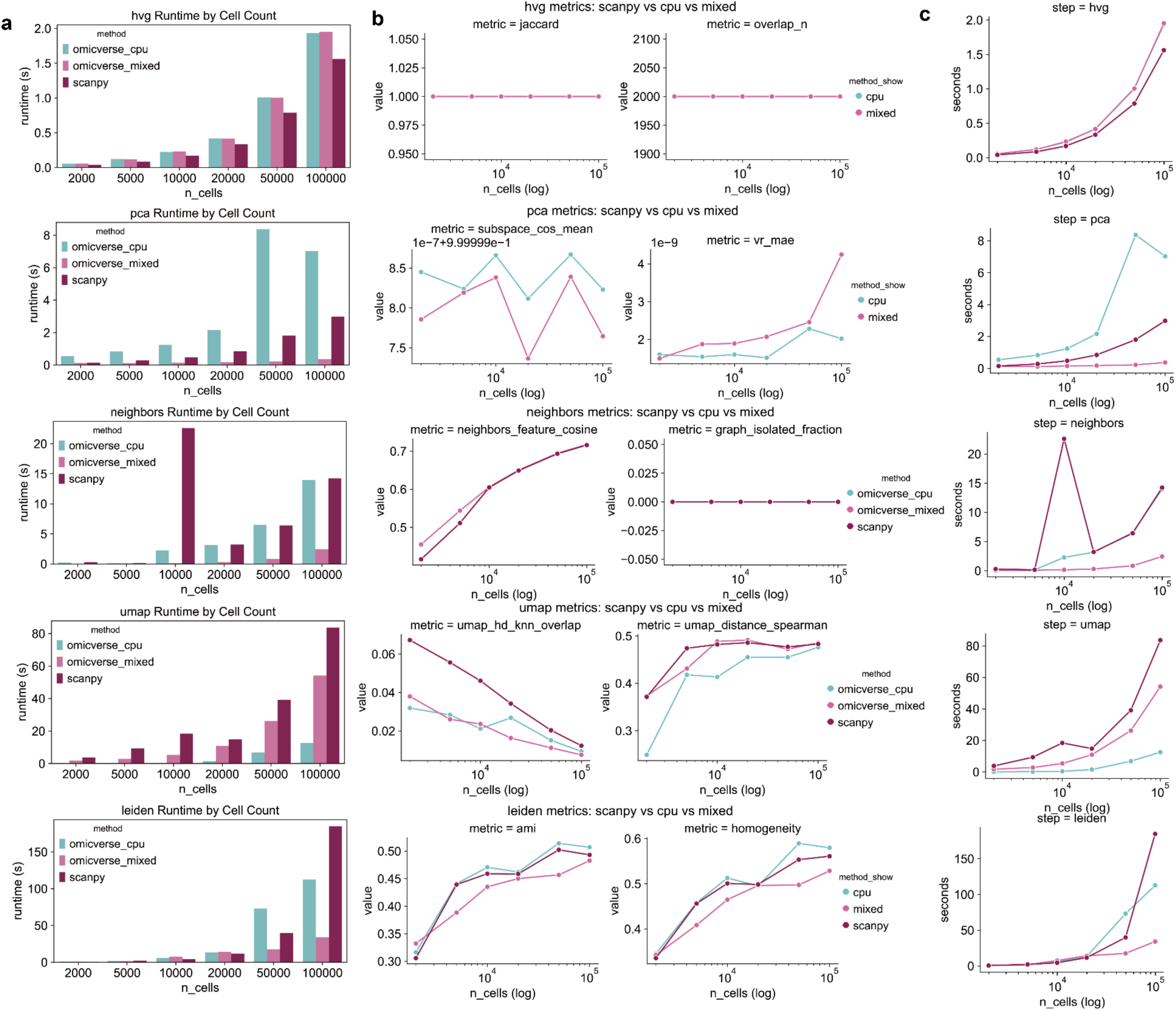
Benchmarking runtime scaling and output fidelity of OmicVerse and Scanpy across increasing cell numbers. **(A)** Stepwise runtime comparison for highly variable gene selection (HVG), principal component analysis (PCA), neighbor graph construction, UMAP embedding, and Leiden clustering across datasets ranging from 2,000 to 100,000 cells. Bars show wall-clock time for Scanpy, OmicVerse CPU, and OmicVerse mixed backends. **(B)** Accuracy and structural fidelity metrics for the same benchmark. For HVG, agreement with the Scanpy reference was quantified by the Jaccard index and the number of overlapping selected genes (overlap_n). For PCA, agreement with the Scanpy reference was measured by mean subspace cosine similarity (subspace_cos_mean) and mean absolute error of explained variance ratios (vr_mae). For the neighbor graph, intrinsic graph quality was assessed by the mean cosine similarity between each cell and its graph neighbors in high-dimensional feature space (neighbors_feature_cosine) and by the fraction of isolated nodes (graph_isolated_fraction). For UMAP, embedding faithfulness was evaluated using overlap between high-dimensional and UMAP-space k-nearest neighbors (umap_hd_knn_overlap) and the Spearman correlation between high-dimensional and UMAP pairwise distances (umap_distance_spearman). For Leiden clustering, agreement with ground-truth cell labels was quantified using adjusted mutual information (AMI) and homogeneity. **(C)** Runtime growth curves for each major analysis step as a function of cell number, plotted on a logarithmic x axis. Together, these analyses compare both computational efficiency and biological/structural consistency across implementations.

**Supplementary Figure 2.**
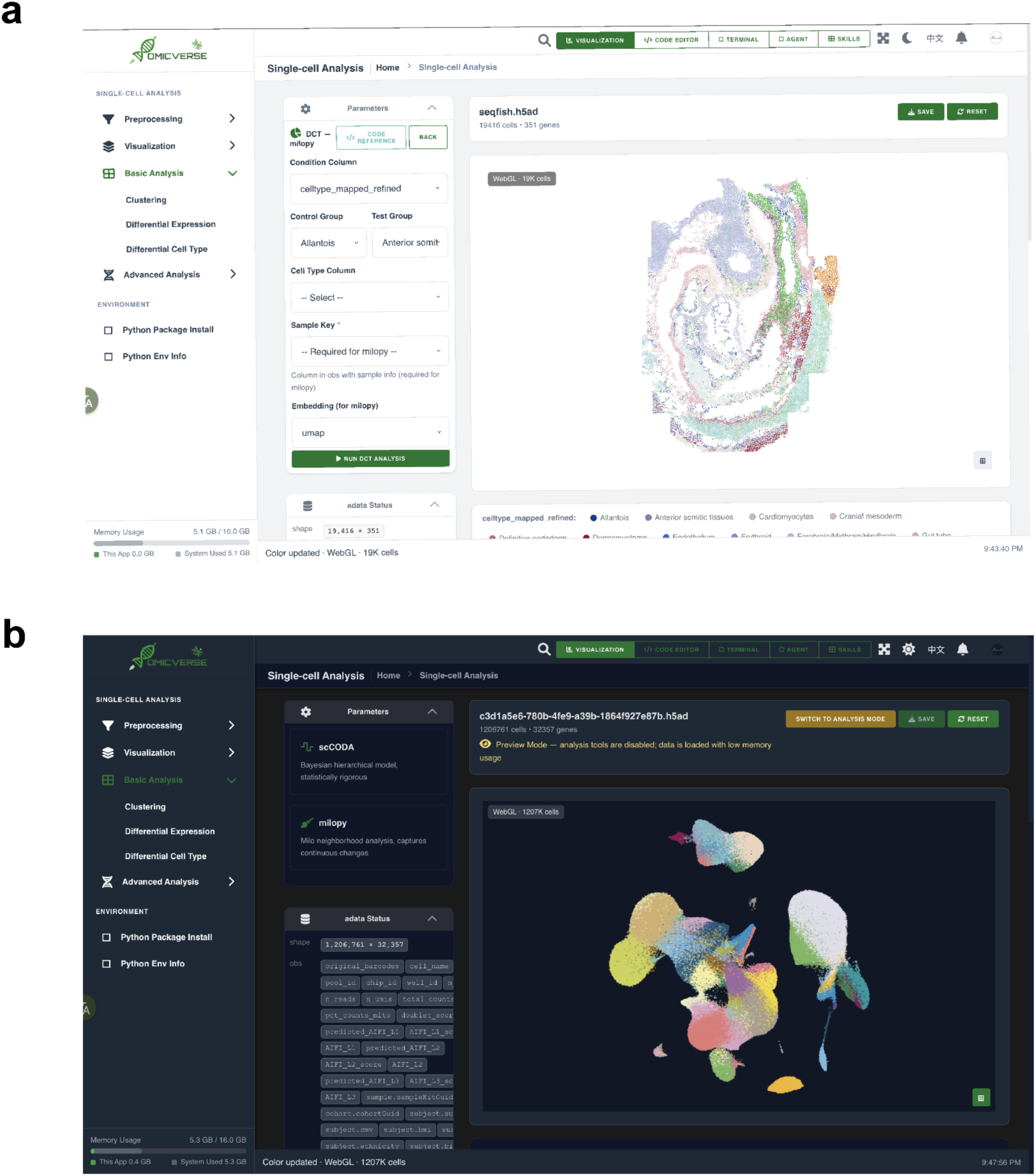
Visualization workspace for scalable single-cell and spatial data exploration. **(A)** Representative view of the OmicVerse Web visualization workspace in analysis mode, showing an interactive single-cell analysis interface with a parameter configuration panel, dataset summary, and embedding-based visualization rendered in the central canvas. The left navigation sidebar organizes preprocessing, visualization, basic analysis, clustering, differential expression, differential cell type analysis, and advanced analysis modules within a unified workflow. **(B)** Representative large-scale dataset view in preview mode, illustrating low-memory loading and WebGL-based rendering for high-cell-count data. In this mode, the interface supports rapid structural inspection of a large .h5ad dataset before switching to full analysis mode, demonstrating that the visualization workspace is designed not only for routine small- to medium-scale exploration but also for interactive inspection of atlas-scale single-cell datasets.

**Supplementary Figure 3.**
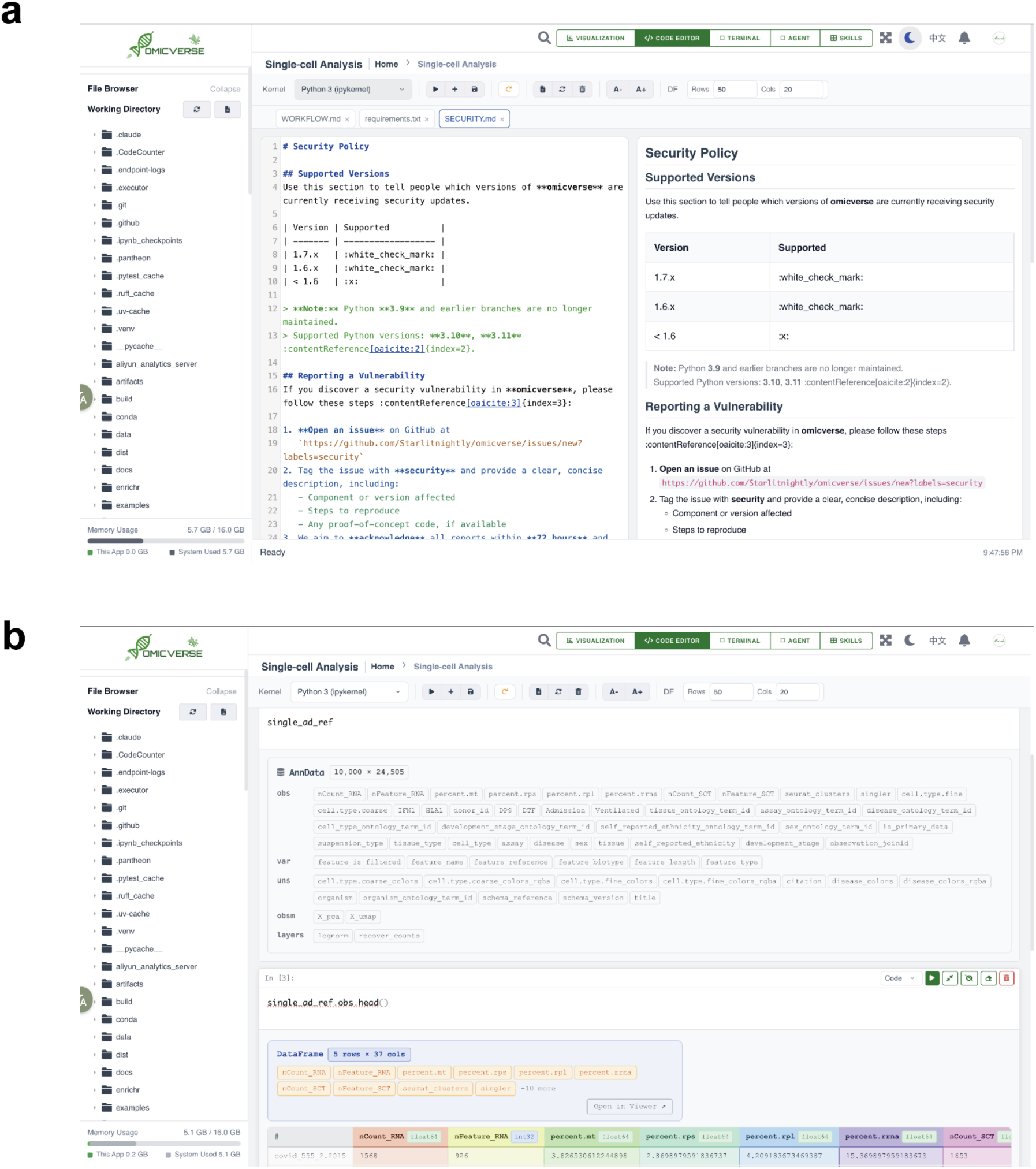
Notebook-style code editor with structured object inspection and live document preview. **(A)** The OmicVerse Web code editor displaying a Markdown document (SECURITY.md) in a split-pane layout, with source editing on the left and live rendered preview on the right. This interface supports notebook-style editing together with direct authoring and preview of workflow documentation and configuration files. **(B)** Structured inspection of biological analysis objects within the notebook environment. An AnnData object is shown as a summarized object card with compact views of obs, var, uns, obsm, and layers, while a DataFrame output is presented as a bounded tabular preview rather than full object expansion. Together, these views illustrate memory-aware inspection of large omics objects in the browser-based execution environment.

**Supplementary Figure 4.**
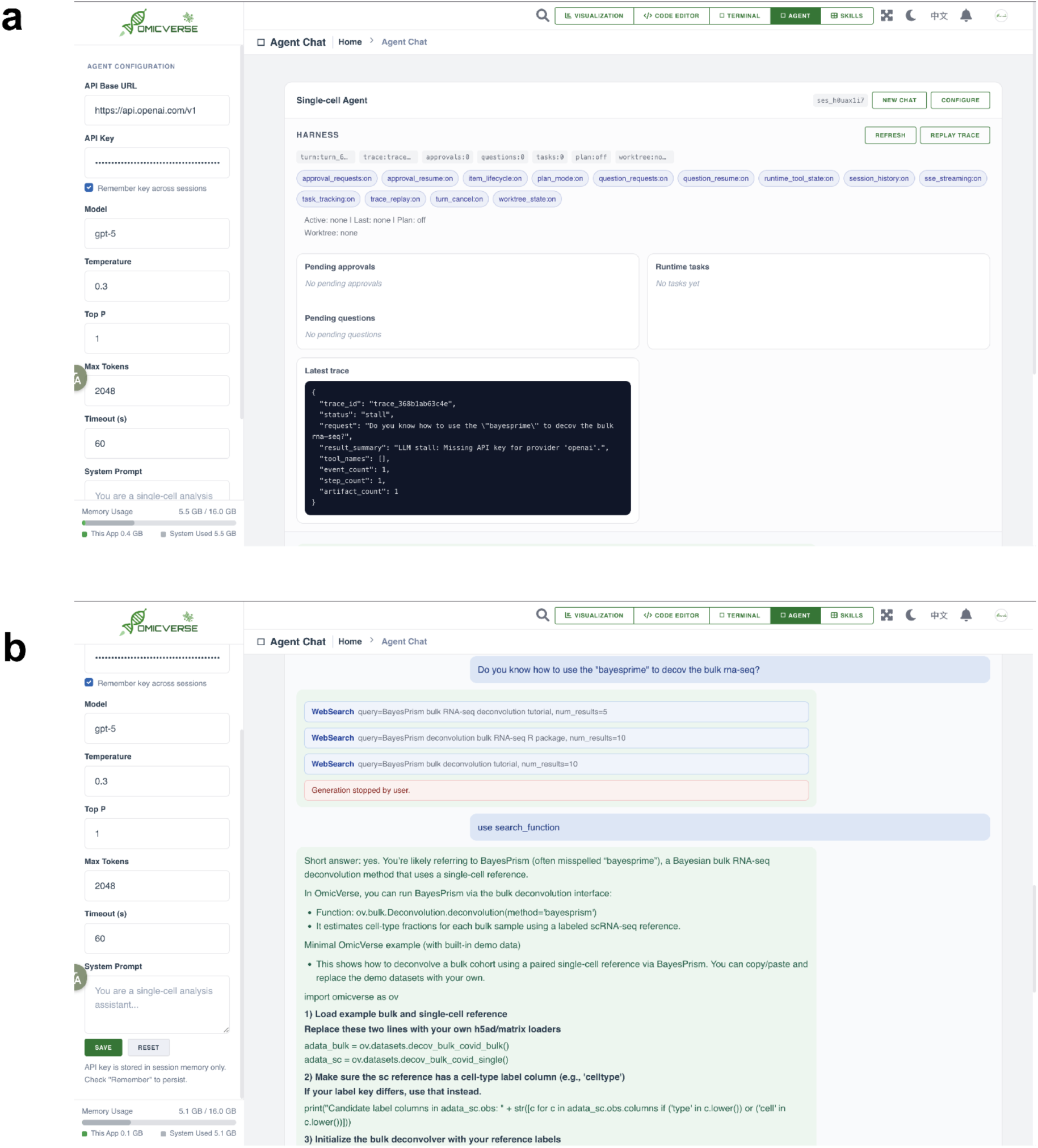
Stateful and auditable agent workspace for human-in-the-loop analysis assistance. **(A)** Agent configuration and runtime overview in the OmicVerse Web agent workspace. The interface exposes model and endpoint settings together with session-level controls and runtime state summaries, including approvals, questions, tasks, plan mode, trace information, and replay controls, thereby making agent execution state visible and inspectable. **(B)** Example multi-turn interaction in the agent chat interface. Tool invocations are displayed inline during execution, and the final response is streamed back into the conversation view. This panel illustrates that the agent workspace is integrated with explicit tool-use tracing and interruption-aware interaction, rather than functioning as an isolated black-box chat window.

**Supplementary Figure 5.**
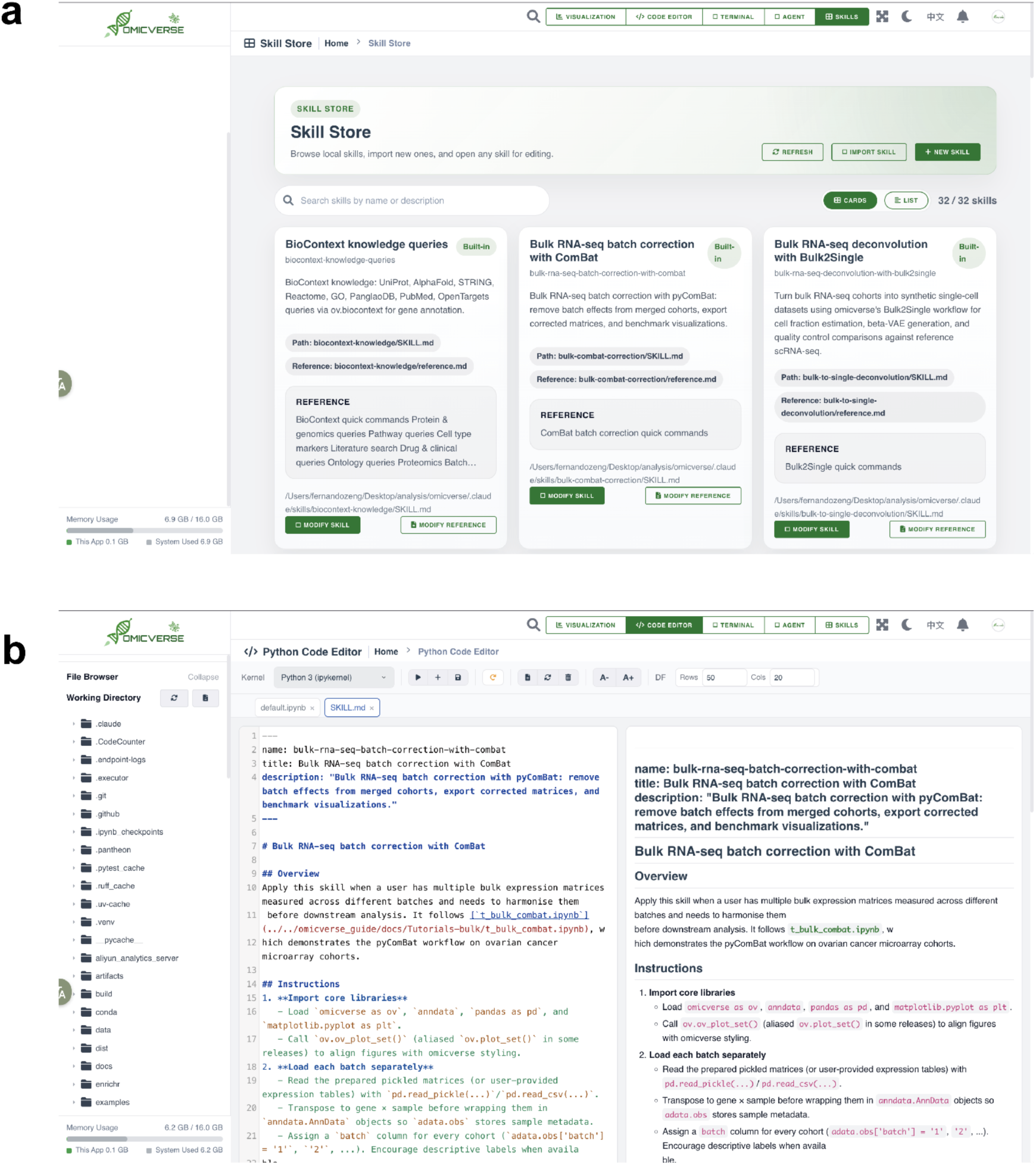
Skill Store and integrated skill editing workflow for reusable analysis knowledge. **(A)** Skill Store interface for managing reusable local skills in OmicVerse Web. The workspace supports browsing, searching, importing, refreshing, and creating skills, while displaying per-skill metadata and associated reference documents in a card-based view. **(B)** Direct editing of a SKILL.md file in the integrated code editor, shown with split-pane source editing and rendered Markdown preview. This design enables users to curate reusable analytical workflows and method notes within the same session in which data analysis is performed, linking skill authoring to downstream agent discovery and workflow reuse.

## Supplementary Table

Supplementary Table 1 | **OmicVerse’s omics analytical methods and alternative implementations in the open-source computational biology ecosystem.** This table summarizes the analytical methods integrated into OmicVerse that are broadly useful for single-cell, spatial, bulk, and multi-omics data analysis, together with implementations of the same or related methods in other open-source packages, when available. Basic data structures, visualization utilities, and general-purpose preprocessing operations are not included unless they represent methodologically distinct analytical modules. The information was updated on 2026.03.11. All listed packages must have publicly accessible source code. The inclusion criteria for “other Python implementations” are: 1) the package is publicly released and installable through standard distribution channels (for example, PyPI or Conda); 2) the repository (GitHub or others) has at least 10 stars; and 3) the package provides an independent implementation of the relevant method(s) rather than simply wrapping OmicVerse functionality. The column “Notable R implementation” provides one representative example for each method instead of an exhaustive survey. We specifically include R implementations because of the long-standing and influential role of the R ecosystem in bioinformatics and omics data analysis.

supp_table_1.xlsx

Supplementary Table 2 | **Registered OmicVerse functions and their corresponding implementations within the open-source omics analysis ecosystem.** This table summarizes the user-facing functions registered in OmicVerse across bulk, single-cell, spatial, preprocessing, plotting, and utility modules. For each function, we provide a brief functional description, a representative calling pattern, GPU support status when available, the corresponding source or registry anchor, and the typical input data structure. The table is intended to document the callable function registry that supports OmicVerse’s unified analytical interface, rather than to provide an exhaustive benchmark of all supported methods. Low-level data structures, internal helper functions, and non-registered backend utilities are not included unless they are exposed as user-accessible functions. The information was updated on 2026.03.11. All listed functions correspond to publicly accessible source code in the OmicVerse repository.

supp_table_2.xlsx

**Supplementary Table 3** | This table lists scientific research papers that used OmicVerse in their analyses. Papers were identified on **March 6, 2026** by using Lens (lens.org), with the **OmicVerse paper Lens ID (050-069-843-432-178)** as the search anchor and an exact search for the keyword **“OmicVerse”**, to obtain a candidate set of scholarly works citing the paper (“citing scholarly works”). To prevent irrelevant records from being included, we manually reviewed each candidate paper and retained only those that explicitly mentioned **OmicVerse/omicverse** as part of their analytical workflow in the **Methods/analysis** sections (e.g., specific module/function calls, workflow descriptions, or algorithmic implementations based on OmicVerse). Studies that cited the OmicVerse paper only in the Discussion or Related Work sections without describing actual use of the OmicVerse tool were excluded. Likewise, studies that used OmicVerse only indirectly through other packages, without explicitly mentioning OmicVerse in the paper, were excluded. For papers confirmed as “actual use”, we further recorded their usage scenarios (e.g., quality control and preprocessing, batch correction, deconvolution, gene set scoring, trajectory inference, enrichment analysis, and spatial analysis) and summarized them in the **“OmicVerse-Analysis”** column. Journal impact factors (IF) were collected from a standardized source (e.g., a specified JCR year) and the table was sorted by impact factor. Citation counts were obtained from Google Scholar on the same date. We acknowledge that this list may miss some papers due to limitations of search engine coverage, version differences (preprint vs. published version), or inconsistencies in citation formats. For transparency, if an “overlap” column is included, a “Y” indicates that the paper’s author list overlaps with the author list of the original OmicVerse paper (or a designated author set specified by our group).

supp_table_3.xlsx

